# Kisspeptin fiber and receptor distribution analysis suggests its potential role in central sensorial processing and behavioral state control

**DOI:** 10.1101/2023.09.05.556375

**Authors:** Limei Zhang, Vito S. Hernández, Mario A. Zetter, Oscar R. Hernández-Pérez, Rafael Hernández-González, Ignacio Camacho-Arroyo, Lee E. Eiden, Robert P. Millar

## Abstract

**Background:** Kisspeptin (KP) signaling in the brain is defined by the anatomical distribution of KP-producing neurons, their fibers, receptors, and connectivity. Technological advances have prompted a re-evaluation of these chemoanatomical aspects, originally studied in the early years after the discovery of KP and its receptor *Kiss1r.* We have previously characterized(1) seven KP neuronal populations in the mouse brain at the mRNA level, including two novel populations, and examined their short-term response to gonadectomy.

**Methods:** In this study, we mapped KP fiber distribution in rats and mice using immunohistochemistry under intact and short- and long-term post-gonadectomy conditions. *Kiss1r* mRNA expression was examined via RNAscope, in relation to vesicular GABA transporter (*Slc32a1*) in whole mouse brain and to KP and vesicular glutamate transporter 2 (*Kiss1* and *Slc17a6*) in hypothalamic RP3V and arcuate regions.

**Results:** We identified KP fibers in 118 brain regions, primarily in extra-hypothalamic areas associated with sensorial processing and behavioral state control. KP-immunoreactive fiber density and distribution were largely unchanged by gonadectomy. *Kiss1r* was expressed prominently in sensorial and state control regions such as septal nuclei, the suprachiasmatic nucleus, locus coeruleus, hippocampal layers, thalamic nuclei, and cerebellar structures. Co-expression of *Kiss1r* and *Kiss1* was observed in hypothalamic neurons, suggesting both autocrine and paracrine KP signaling mechanisms.

**Conclusion:** These findings enhance our understanding of KP signaling beyond reproductive functions, particularly in sensorial and behavioral state regulation. This study opens new avenues for investigating KP’s role in controlling complex physiological processes, including those not related to reproduction.

## 1. Introduction

Kisspeptin (KP) is a neuropeptide encoded by the *Kiss1* gene, initially known as metastin, and identified for its activity in inhibiting cancer metastasis(2). It is now well-recognized as a critical regulator of the mammalian reproductive system, playing a pivotal role in the activation of the hypothalamo-pituitary-gonadal (HPG) axis by stimulating the release of gonadotropin-releasing hormone (GnRH)(3, 4). Beyond its reproductive functions, KP is increasingly implicated in a variety of physiological processes, including the regulation of emotions and cognitive functions(5), especially, although not limited to, those involved in sexual behavior(6). For instance, kisspeptin receptors are expressed in the amygdala and hippocampus, critical regions for emotional processing, and have been associated with anxiolytic/anxiogenic effects and the modulation of stress(7–9) and fear responses(10–12). Additionally, KP has been implicated in cognitive functions such as learning and memory, potentially through its actions within the hippocampus and prefrontal cortex(13). Thus, KP is likely to play a multifaceted role in brain function, beyond control of the HPG axis.

In a recent study in mouse, our group reported a comprehensive KP cell population description and molecular signature characterization within the excitatory/inhibitory (glutamatergic-GABAergic) context throughout mouse brain, using a sensitive dual and multiple channel in situ hybridization method. We described chemotypes of seven distinct KP populations, four of them within the hypothalamus and three of them as extrahypothalamic populations. The KP cell groups located in the ventral division of the premammillary nucleus of the hypothalamus and the nucleus of tractus solitarii in the brain stem, are new additions to the literature. Their up- and down regulation by short-term gonadectomy (GNX) was also reported quantitatively(1). These results prompted us to extend the study by focusing on kisspeptin immunopositive fiber distribution in both rat and mouse, and on rodent KP receptor (*Kiss1r*) distribution using the RNAscope method.

There had been comprehensive anatomical studies on KP neuronal populations and fiber distributions in the brain of diverse animal species in the literature of the first decade after the discovery of KP, albeit hypothalamic, rather than extrahypothalamic regions, were the main focus of those studies due to the role of KP in HPG axis regulation(13–18). However, KP may also regulate sexual behavior, in additional to reproductive endocrinology, within the mammalian brain(6). Due to the emerging evidence for kisspeptin function beyond its well-documented role in orchestration of GnRH secretion in the hypothalamus, more comprehensive data on the distribution of kisspeptin cell bodies and nerve fibers throughout the brain are needed. In particular, some data have been questioned as potentially unreliable, due to possible adventitious cross-reactivity with other RFamide-peptides(16). We have used a KP antibody with well-documented authenticity and lack of cross-reactivity, and applied state-of-the-art tissue preparation and staining methods (19) in this study. Providing a reliable and comprehensive description of the distribution of KP cell bodies and fibers throughout the brain should contribute to a better understanding of the possible role(s) of KP first messenger function relevant to both reproductive and non-reproductive behavior. We report here 118 brain regions that contain KP fibers and terminals, with most of them located in extra-hypothalamic regions closely involved in central sensorial processing and behavioral state control(20–25). KP fiber density was similar in long-term (one year) gonadectomized compared to intact male or female mice.

The kisspeptin receptor (Kiss1R), was first identified in rat in 1999 as an orphan receptor previously called Gpr54(26). Kiss1R is a metabotropic receptor coupled to Gq/11, and its inactivation has been associated with hypogonadism(27). After KP binds its receptor, phospholipase C (PLC) is activated and hydrolyzes phosphatidylinositol 4,5-bisphosphate (PIP2) to produce diacylglycerol (DAG) and inositol 1,4,5-trisphosphate (IP3) with a rise in Ca++ mobilized from intracellular stores and the activation of protein kinase C (PKC)(28, 29). There have been few reports on Kiss1R expression in the CNS(27, 28, 30, 31) and a comprehensive description, especially Kiss1R co-expression within the context of excitatory and inhibitory neurotransmission, is incomplete. In this study, we used the RNAscope dual and multiplex in situ hybridization method to map the Kiss1r expression throughout mouse brain. *Kiss1r* was prominently expressed in brain sensory and state control regions, such as main and accessory olfactory bulbs, septal nuclei, tuberomammillary nuclei, the suprachiasmatic nucleus, the locus coeruleus, the hippocampal pyramidal and granule cell layers, the thalamic nuclei, and the cerebellar granule cell layer, as well as the interposed nucleus. *Kiss1r* was also co-expressed in *Kiss1*-expressing hypothalamic neurons and neighboring cells, suggesting the KP system uses autocrine and paracrine mechanisms. Data from this study provide additional anatomical features of the KP system for further hypothesis generation and testing of the role of KP beyond HPG axis regulation.

## 2. Material and methods

### 2.1. Animals

Wistar rats (N=12, male = 6, female = 6) and C57BL/6 mice (N=30, male = 15, female = 15) were obtained from the local animal vivarium and housed three per cage under controlled temperature and illumination (12h/12h) with water and food *ad libitum*. Mice subjected to gonadectomy were singly housed for recovery for one week after surgery and then returned to their home cages. All animal procedures were approved by the local research ethics supervision commissions (license number CIEFM-079-2020, UNAM and NIMH/ACUC approval LCMR-08).

To evaluate the effect of gonadectomy on the expression of kisspeptin throughout the brain in male and female mice, we compared the KP immunoactivity of 24 mice distributed in the following experimental groups:

| Male |  |  |  | Female |  |  |  |
| --- | --- | --- | --- | --- | --- | --- | --- |
| Five month old |  | Fifteen month old |  | Five month old |  | Fifteen month old |  |
| Intact | 1 mo ORX | Intact | 12 mo ORX | Intact | 1 mo OVX | Intact | 12 mo OVX |
| n=3 | n=3 | n=3 | n=3 | n=3 | n=3 | n=3 | n=3 |

For RNAscope ISH experiments, we used three male and three female mice of three months of age. Vaginal cytology was performed in female mice to assess the estrous cycle. Brains were collected during the proestrus stage associated with a high level of estrogen.

### 2.2. Gonadectomy

#### 2.2.1 Mouse ovariectomy (OVX)

A dorsal approach to remove the ovaries was used (32). Briefly, animals were anaesthetized with ketamine 90 mg/kg + xylazine 20 mg/kg *i.p.* Under anesthesia, animals were placed in ventral recumbency and a 3 cm/side square region centered over the third lumbar vertebra shaved. The shaved area was disinfected with Povidone and 70% alcohol, and a sterile fenestrate drape was placed over it. A single midline dorsal incision approximately 1 cm long was made in the skin at the level of the bottom of the rib cage. The incision was moved laterally to cut the right and left abdominal walls. After cutting the wall (oblique and transverse abdominal muscles), blunt dissection through the body wall with mosquito hemostats was performed. Forceps were gently and carefully introduced to retrieve and exteriorize the ovary. The ovary, ovarian vessels, and ligament were ligated with 4-0 synthetic absorbable sutures and cut. The wall muscle layer incision was closed with a 4-0 synthetic, monofilament, absorbable suture. The skin incision was closed with a subcuticular suture.

#### 2.2.2 Mouse orchiectomy (ORX)

For the excision of the testicles, we followed a midline incision technique(33). Briefly, animals were anaesthetized with ketamine 75 mg/kg + xylazine 16 mg/kg i.p. and placed on the operating table on their backs. The central ventral side of the scrotum area was disinfected with Povidone and 70% alcohol. A single incision on the ventral side of the scrotum (1 cm long) penetrating the skin was performed. Each cremaster muscle was cut, and each testis was exposed by gently pulling it through the incision using blunt forceps. Cauda of the epididymis, caput epididymis, vas deferens, and testicular blood vessels were exposed while holding the testicular sac with sterile tooth forceps. A single ligature 4-0 synthetic absorbable suture around the vas deferens and blood vessels was made, prior to excision and gentle removal of the testis. The remaining content was replaced in each one of the testicular sacs. The skin incision was closed with a subcuticular suture.

### 2.3 Immunohistochemistry (IHC)

Animals were deeply anaesthetized with sodium pentobarbital (100 mg/kg, *i.p.*). They were perfused transaortically with 0.9% saline followed by a cold fixative containing 4% paraformaldehyde in 0.1 M sodium phosphate buffer (PB, pH 7.4) plus 15% v/v saturated picric acid for 15 min. Brain tissues were rinsed immediately after the 15 min fixation, until the buffer was colorless. Brains were then sectioned the same day, using a Leica VT 1000S vibratome at 70 μm thickness, in the coronal and parasagittal planes, and the immunohistochemical (IHC) reaction was performed the same day. This procedure yielded a more comprehensive IHC labelling than longer fixation and delayed IHC as previously reported and discussed(19). One of every three sections was selected, and sections from animals that would be compared (*i.e*., female control *vs.* female gonadectomy) were incubated in the same vial. To block the unspecific antibody binding, sections were incubated for one hour in a solution consisting of 20% normal donkey serum (NDS) diluted in Tris-buffered saline (TBS, pH 7.4) plus 0.3% Triton-X (TBST). After the blocking step, sections were incubated for 48 h at 4°C in a polyclonal sheep anti-kisspeptin antibody(34) (GQ2, kindly supplied by Mohammad Ghatei†, Imperial College, London, UK), diluted at a 1:5000 concentration in TBST + 1% NDS. This antibody recognizes kisspeptins 54, 14, 13, and 10 that conserve the C-terminal decapeptide essential for biological activity and shows virtually no cross-reactivity (<0.01%) with other related human RF-amide peptides(34). After three wash steps, sections were incubated for 24 h with the secondary donkey anti-sheep biotinylated antibody (Cat: 713-065-147, Jackson ImmunoResearch Inc, PA, USA) diluted at a concentration of 1:500 in TBST + 1% NDS. To detect the biotinylated antibody, VECTASTAIN Elite ABC HRP Kit (Cat: PK-6100, Vector Laboratories, Inc. CA, USA) was used, with H_2_O_2_ and 3,3’-diaminobenzidine as substrates. Finally, sections were mounted on glass slides, dehydrated, dipped in xylene, and cover-slipped using Permount^TM^ mounting medium.

### 2.4 Anatomical assessment of kisspeptin innervation density and comparison between control and gonadectomized animals

To assess the projection patterns of kisspeptin immunopositive fibers throughout the brain four researchers independently examined the slides under a light microscope, after identifying the regions based on the Paxinos Rat & Mouse brain atlases(35, 36). Scores of "+" (sparse, main axons passing, without branching nor axon terminals); "++", scattered, some branching; "+++", moderate, abundant branching and axon terminals; "++++", intense, extensive branching and intense axon terminals, with confluent fibers that it was difficult to see a separation between them, were reported.

### 2.5 Anatomical assessment of *Kiss1r* mRNA expression in the context of Slc32a1-expressing neurons throughout the brain

Six C57Bl mice, 3 male and 3 female of 12-14 weeks old, were deeply anesthetized with sodium pentobarbital (100 mg/kg b.w., i.p.) and decapitated using a small animal guillotine. The brains were then extracted and rapidly frozen in Dry Ice. Fresh-frozen brains were sectioned sagittally into 12 μm thick slices using a Leica CM-1520 cryostat and mounted onto positively charged Superfrost Plus microscope slides (Cat. 22-037-246, Fisher Scientific, Pittsburgh, PA). The detailed methods have been previously described (1, 37, 38).

Probes used for the detection of mRNAs in the double or multi-channel *in situ* hybridization (DISH and MISH) were custom-designed by Advanced Cell Diagnostics (https://acdbio.com/rnascope-25-hd-duplex-assay) for kisspeptin (Mm-Kiss1, channel 1 probe), VGAT (Mm-Slc32a1, channel 1 and channel 4 probes), kisspeptin receptor (Mm-Kiss1r, channel 2 probe) and VGLUT2 (mm-*Slc17a6*, channel 3). All experimental procedures were carried out in accordance with the manufacturer’s protocol. After sectioning, the slides were fixed in chilled 4% paraformaldehyde (Cat. 30525-89-4, Sigma-Aldrich) prepared in phosphate-buffered saline (Cat. M32631, Sigma-Aldrich), dehydrated in ethanol, and treated with hydrogen peroxide for 15 minutes. They were then treated with protease III for 20 minutes at room temperature, incubated for two hours with the probe mixtures, and underwent amplification and detection steps following the manufacturer’s guidelines. Finally, the slides were counterstained with hematoxylin in case of DISH and cover-slipped.

The processed sections were examined using a Nikon Eclipse E600 microscope along with its digital camera for photographic documentation. Multichannel in situ hybridization (ISH) sections were analyzed using the Stellaris Confocal Microscope from Leica Microsystems. Sections were subsequently scanned using the ZEISS Axio ScanZ.1.

For the semi-quantitative analysis, we determined the percentage of *Kiss1r*-labeled cells that co-expressed *Slc32a1* for the column titled ’Slc32a1 co-expression’ and the total number of Nissl-labeled nuclei (NLN) versus Kiss1r on NLN without Slc32a1 labeling for the column titled ’No Slc32a1 co-expression’ for each region of interest. The number of "+" assigned was based on the following criteria: if the percentage fell within the ranges of 1% to 20% it was assigned (+); from 21% to 40%, (++); from 41% to 60%, (+++); from 61% to 80%, (++++); and from 81% to 100%, (+++++).

## 3. Results and focused discussion

### 3.1 Regional distribution of kisspeptin-expressing fibers and neurons in the mouse and rat brain

In Table 1, we report 118 regions of the rodent brain where we observed kisspeptin (KP immunoreactivity (ir) with varying strengths. We identified only one previous anatomical mapping study on the distribution of KP immunoreactivity in the rat brain that had a similar level of completeness compared to our data. In that study, the authors utilized an antibody produced by Phoenix Pharmaceuticals (39), which has been noted as potentially cross-reactive with other RFamide-related peptides(16). We included the data reported by Brailoiu et al.(39) in rat, using the Phoenix Pharmaceuticals antibody anti-metastin (MT), in the Table 1, highlighted in blue to facilitate comparison with our current findings. All the regions reported in this earlier study were replicated in our experiments, which suggests that the value of the earlier report may have been underestimated. Furthermore, we identified numerous additional regions containing KP-expressing fibers.

**Table 1.**
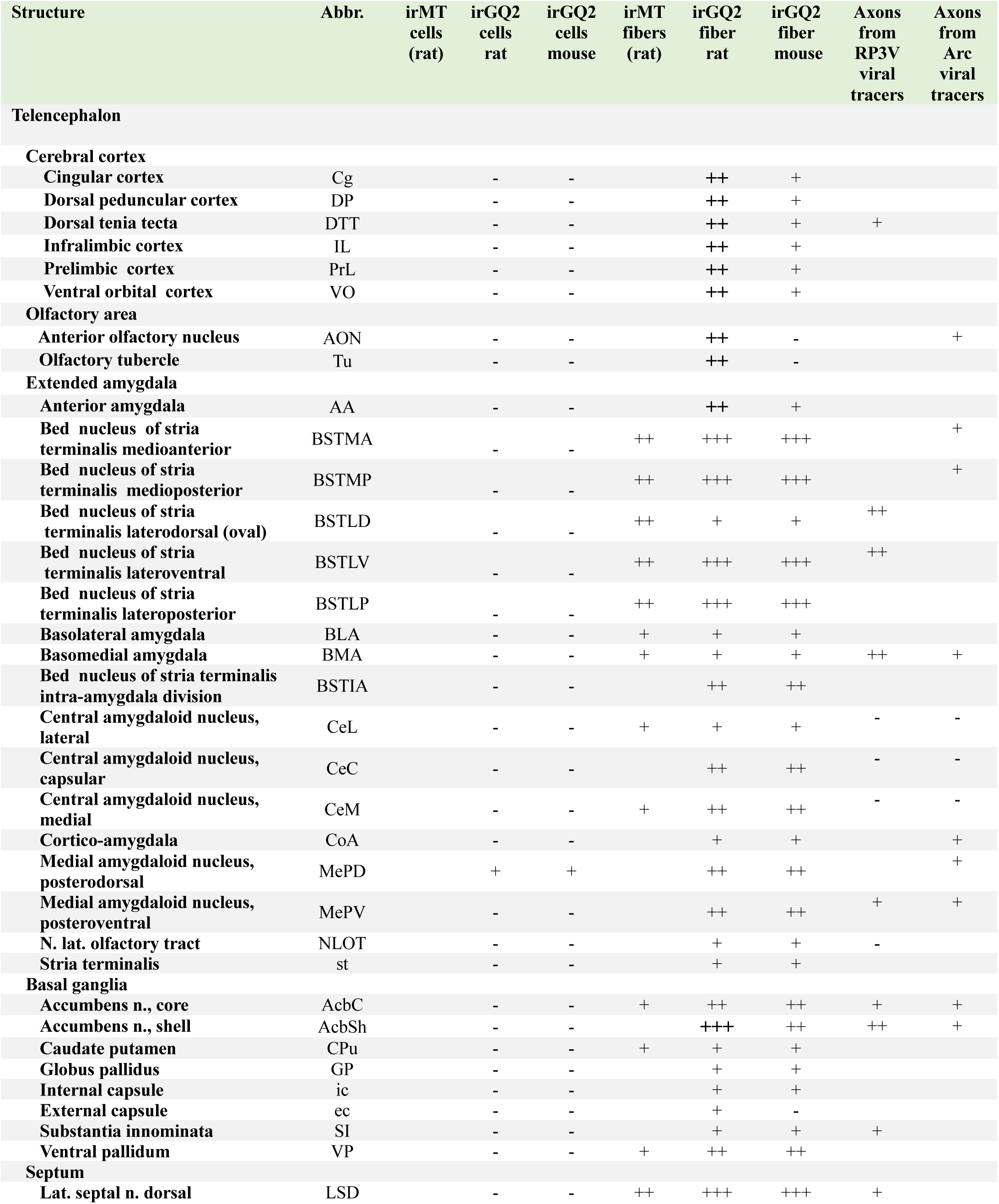
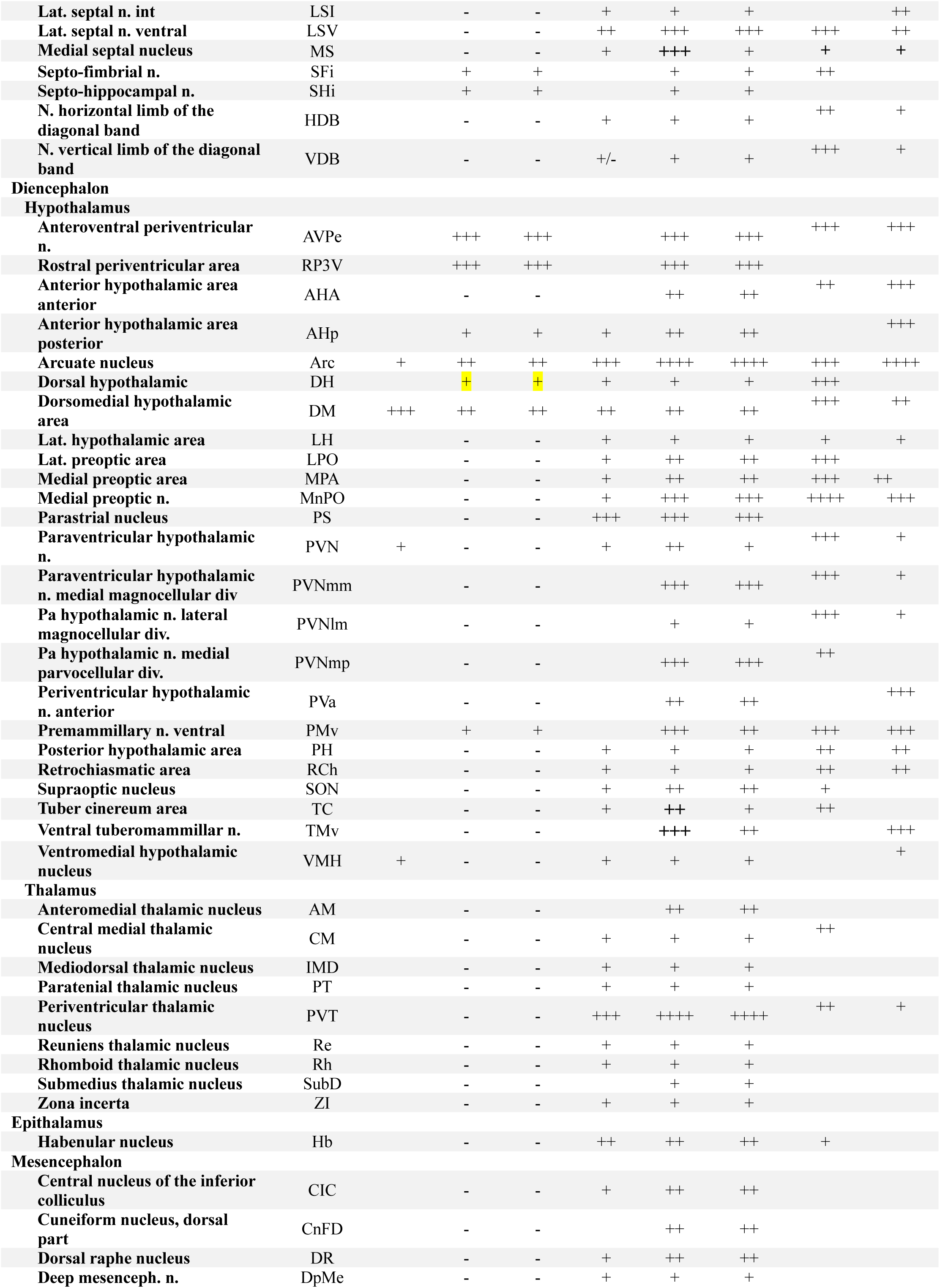
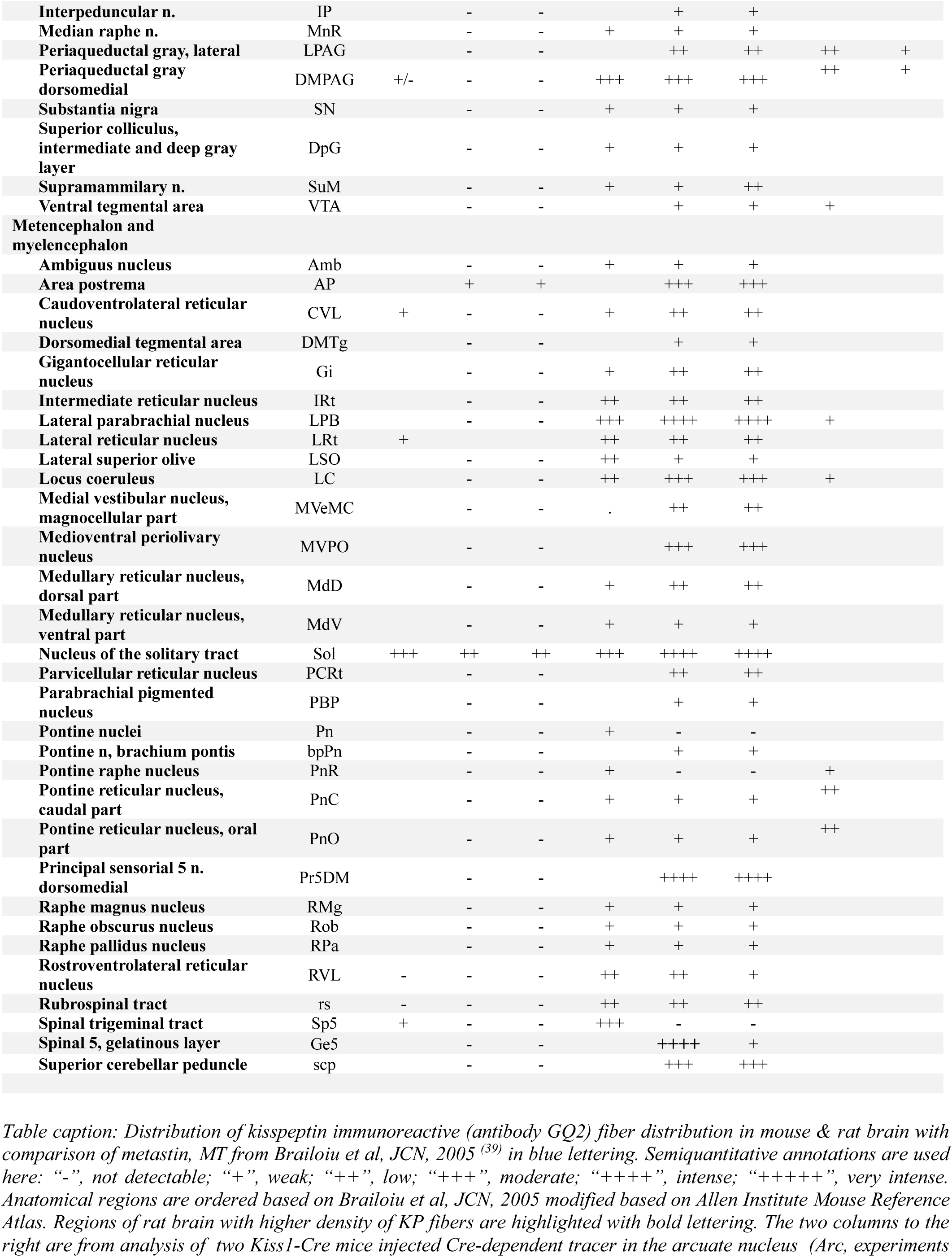
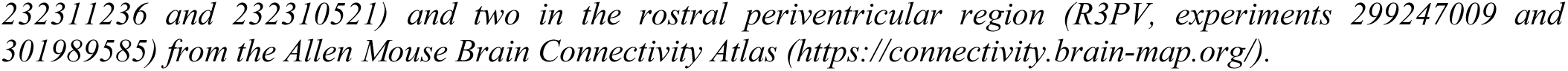
Comparison of the distribution of kisspeptin immunoreactive fibers in mouse and rat brain using antibody GQ2 (current study) to earlier reports using metastin MT antibody (Brailoiu et al, JCN, 2005)

The rat brain, particularly in telencephalic regions (*vide infra*), exhibited stronger immunohistochemical labeling for KP than the mouse brain. However, quantitative conclusions cannot be drawn confidently due to variability among experimental subjects and immunoreactions. Sex differences in the distribution of cells or fibers were inconsistent among experimental subjects, even under identical experimental procedures. For example, within a given region, some reactions were stronger than others, and these differences were not consistently associated with sex or treatment of the subjects. Various unknown factors may have influenced these observations. It is important to note that KP expression may be up- or down-regulated in individual organisms due to their physiological regulation, affecting the presence or absence of peptide vesicles at a given location at the time of perfusion-fixation. Specifically, a kisspeptinergic cell (referring to a neuron that can synthesize Kp and transport it to its axon or dendrite) can only be detected with KP-IHC if the segment is filled with peptidergic vesicles at the moment of fixation. However, determining these factors falls outside the objective of our study, which aims to qualitatively determine the Kp signaling beyond reproductive functions, especially in the brain regions implied to sensory and behavioral states. Based on these observations and rationale, we pooled data from male and female subjects for Table 1 and Figures 1-3, while separating the observations by rodent species (i.e., mouse in Figures 1 and 2 versus rat in Figure 3) to depict the highest possible KP expression in a given region as representative of “ideal subjects.” This scoring reflects the maximum density observed for each region, providing insight into the maximum potential expression of KP in living rodents. In Supplementary Information (SI) Figure 1, we present examples of KP IHC labeling in young male and female mice, both intact and gonadectomized, to further illustrate this point.

**Fig. 1.**
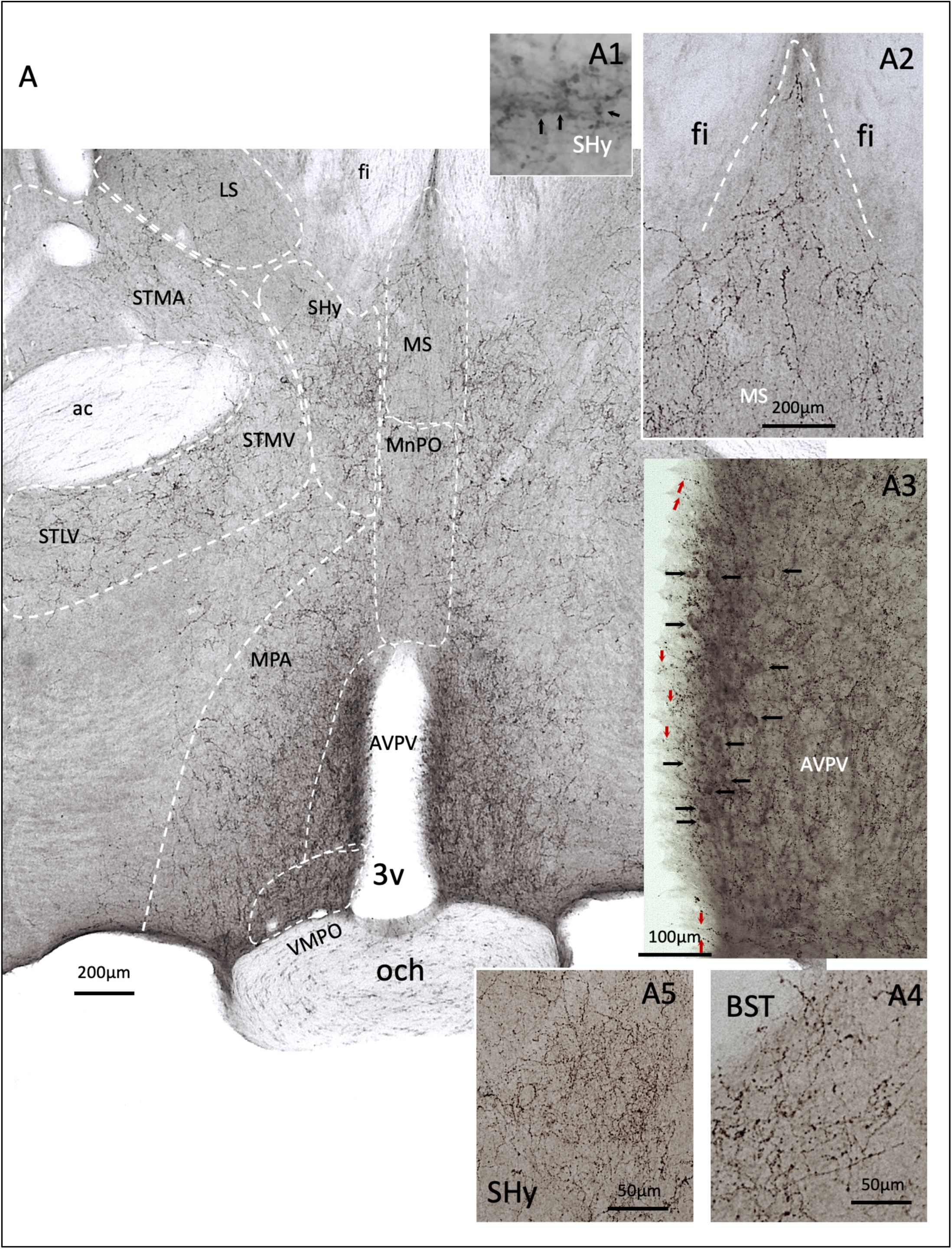
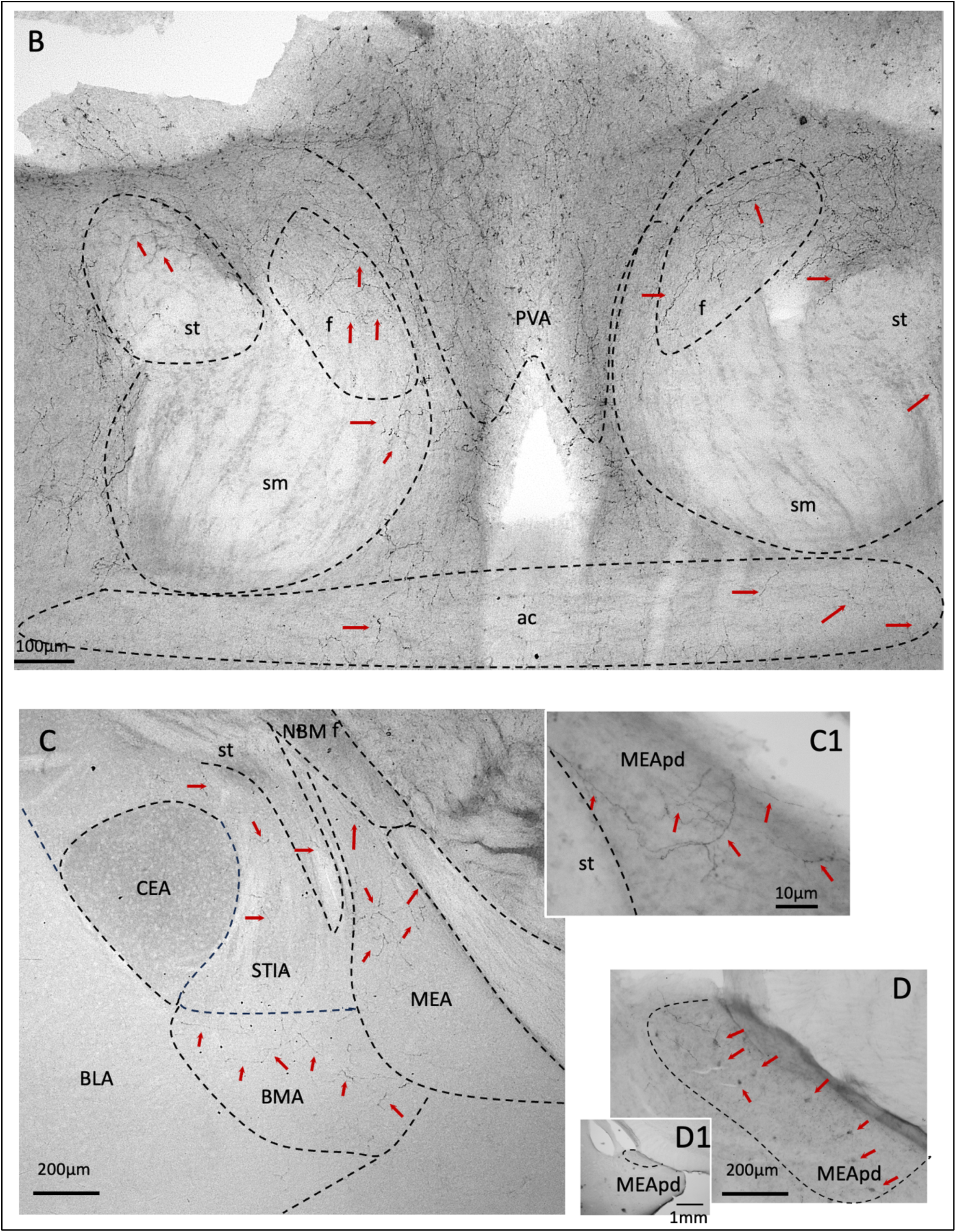
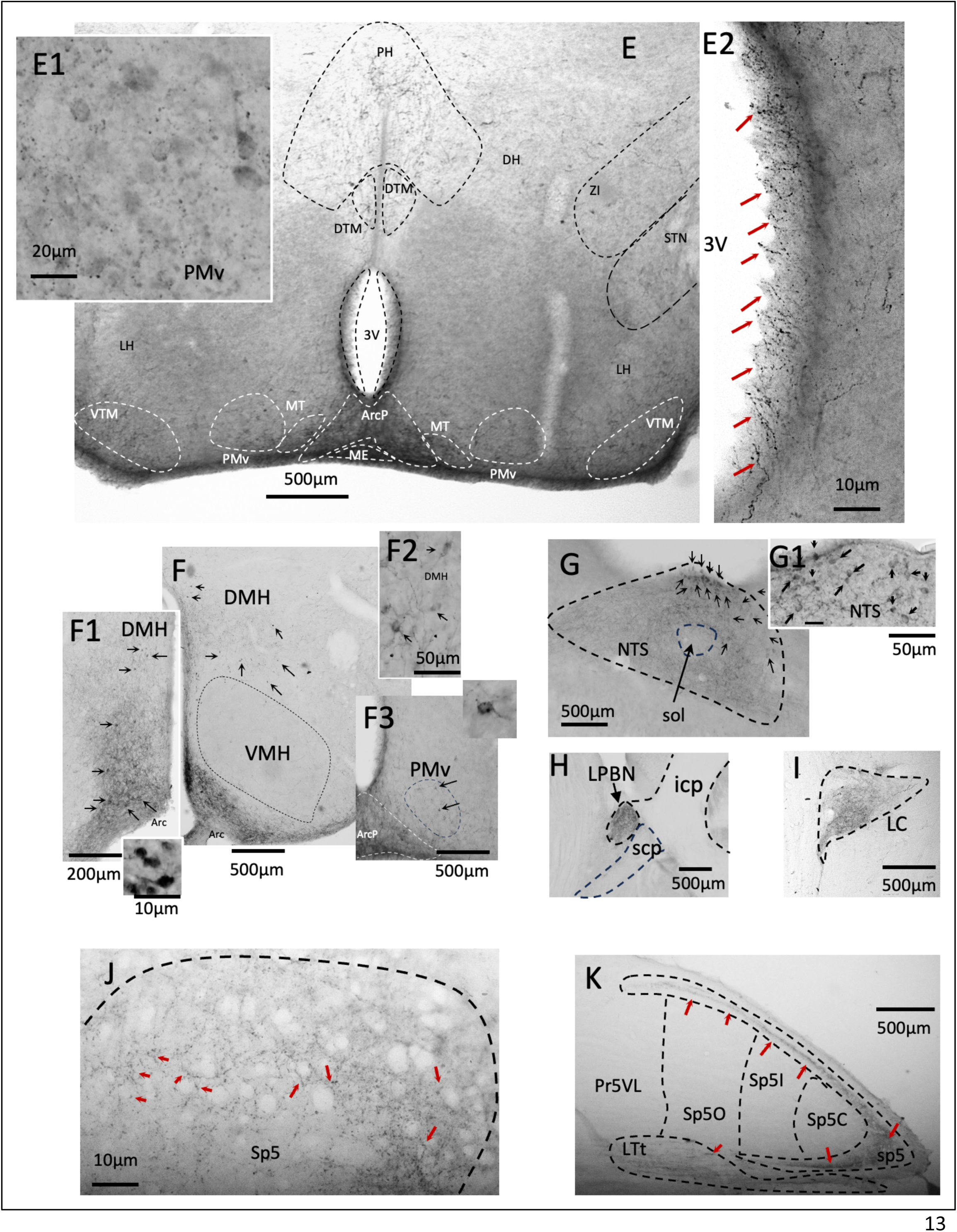
Mouse KP immunohistochemical reactive (-ir) distribution in controlling GnRH secretion, in comparison with other brain regions, with equally intense KP innervation, that are relevant in regulation of brain state. Panel A shows a low magnification micrograph at coronal plane (Bregma 0.38mm dorsal, 0.02mm ventral (35)), where hypothalamic kisspeptinergic regions (regions containing KP-ir cell bodies, such as the antero-ventral periventricular nucleus (AVPV,A3), ventromedial preoptic nucleus (VMPO) are shown. Besides, regions with high densities of KP innervation, which are relevant for brain state control, such as medial nucleus of preoptic area (MnPO), septo-hypothalamic nucleus (SHy, A1, A5, which also contains KP cell bodies, indicated with black arrows. For ISH image of SHy Kiss1 expressing neurons please see (1)), medial septal nucleus (MS, A2)), lateral Septal nu. (LS), bed nucleus of stria terminalis (BST, A4), with its subdivisions medio-anterior (STMA), medio-ventral (STMV) and latero- ventral (STLV). A3 shows the KP-ir in AVPV, where KP-ir neurons are indicated with black arrows and abundant KP-ir fibers innervating the ventricular surface are indicated with red arrows. Panel B, micrograph of a coronal section around Bregma -0.15mm, where 4 brain main conducting fiber systems can be clearly seen, i.e. the anterior commissure (ac), the stria medullaris (sm), the stria terminalis (st) and the fornix (f). KP-ir fibers running inside these conducting systems are indicated with red arrows. Also, the dense innervation pattern of KP-ir fibers in the paraventricular anterior nucleus of the thalamus (PVA) is shown. Panel C shown a low magnification coronal micrograph of amygdala around the coordinate Bregma -1.58mm. KP-ir fibers can be seen in stria terminalis (st), bed nucleus of stria terminalis, intra-amygdalar division (STIA), baso-medial amygdala (BMA), nu. basalis of Meynert (NBM) and medial amygdala (MEA). Interestingly, some fibers within the medial amygdala, postero-dorsal division (MEApd), are observed coming from the st, that the main axon entered the MEApd and branched locally (C1, red arrows). Panel D shows a micrograph of MEApd at Bregma -2.18mm. Numerous KP-ir cell bodies are indicated with red arrows. D1 shows a low magnification micrograph of the panel D. Panels Es and Fs show the microphotographs taken from the hypothalamic tuberal region, where the KP- ir cell bodies are seen in arcuate nucleus (Arc. E and F1), in premammillary nucleus, ventral division (PMv, E and E1, F3) and in the dorso-medial hypothalamic region (DMH, F, F1 and F2, black arrows). KP-ir fibers are observed in other regions relevant for behavioral state control at this level, such as ventral tuberomammillar nucleus (VTM), magnocelular tuberomammillar nucleus (MT) and dorsal tubero mammillary nucleus (DTM), posterior hypothalamus (PH), zona incerta (ZI) and subthalamic nu. (STN) (E). E2 shows the third ventricle at the level of Arc, posterior division (ArcP). Abundant KP-ir fibers extended toward the lumen of the ventricle (red arrows,. E2). Panel Gs- K are micrographs of samples from parasagittal sections of mouse hindbrain. Gs show the KP-ir cell bodies and dense KP-ir fibers in the solitary tract (sol) nucleus (NTS) of mouse medulla. H and I show the dense innervation pattern of KP-ir fibers in pontine nuclei: lateral parabrachial nucleus (LPBN) and locus coeruleus (LC) respectively. J and K show the dense KP-ir fibers in the spinal trigeminal tract (Sp5, K) and in the gelatinous layer of Sp5 (Ge5). Note that within the Ge5, the KP-ir fiber make perisomatic contacts with the cell bodies of the Ge5 (J, red arrows).

**Fig. 2.**
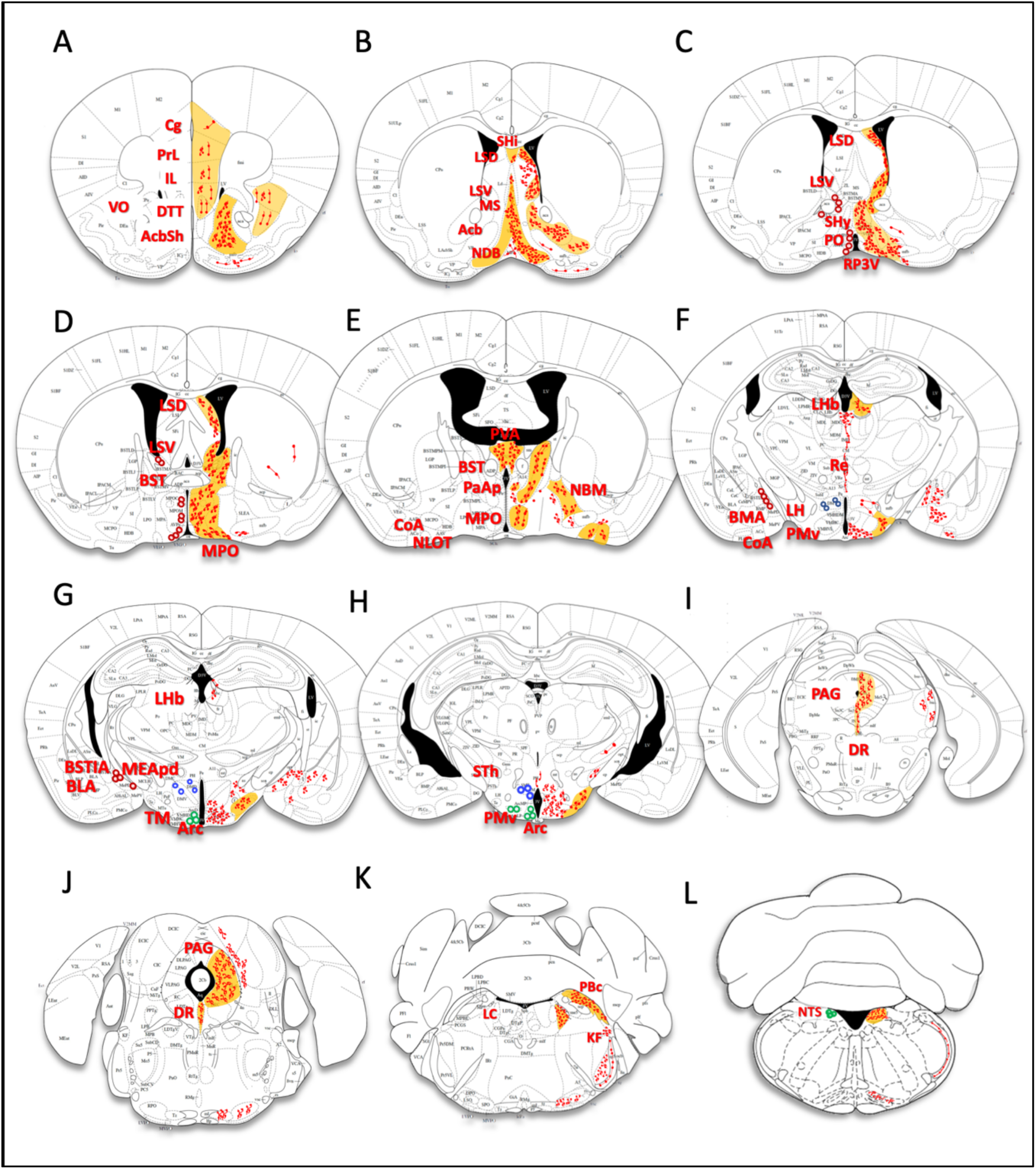
Charting to represent the distribution of kisspeptin immunoreactivity in mouse brain in register with a previous study (14) for purposes of comparison. Sequential atlas planes show in the left side, KP immunoreactive cell bodies from the seven KP cell populations. Color-filled circles symbolizing the molecular signature previously reported (1): KP- glutamatergic (green), KP-GABAergic (mainly, red) and expressing both VGLUT2 and VGAT (blue). The distribution of KP-ir fibers are represented by red lines in the right hemisphere (the thickness of the lines has been exaggerated to ensure that the lines can be seen on the atlas planes). KP-ir fibers observed in brain areas relevant for behavioral state control are shaded in yellow: accumbens nucleus, shell (AcbSh), septohippocampal nu. (SHi), lateral septal nu. dorsal (LSD), medial septal nu. (MS), horizontal division of nucleus of dianogal band of Broca (HDB), septohypothalamic nucleus (SHy), bed nucleus of stria terminalis (BST), medial preoptic area (MPO), paraventricular anterior parvicellular division (PaAP), nucleus basalis of Meynert (NBM), lateral habenula (LHb), lateral hypothalamus (LH), tuberomammillary nu. (TM), ventral premammillary nu. (PMv), periaqueductal grey (PAG), dorsal raphe nu. (DR), locus coeruleus (LC), lateral parabrachial nu. (LPB), nu. of solitary tract (NTS). See Table 1 for remainder of abbreviations.

**Fig. 3.**
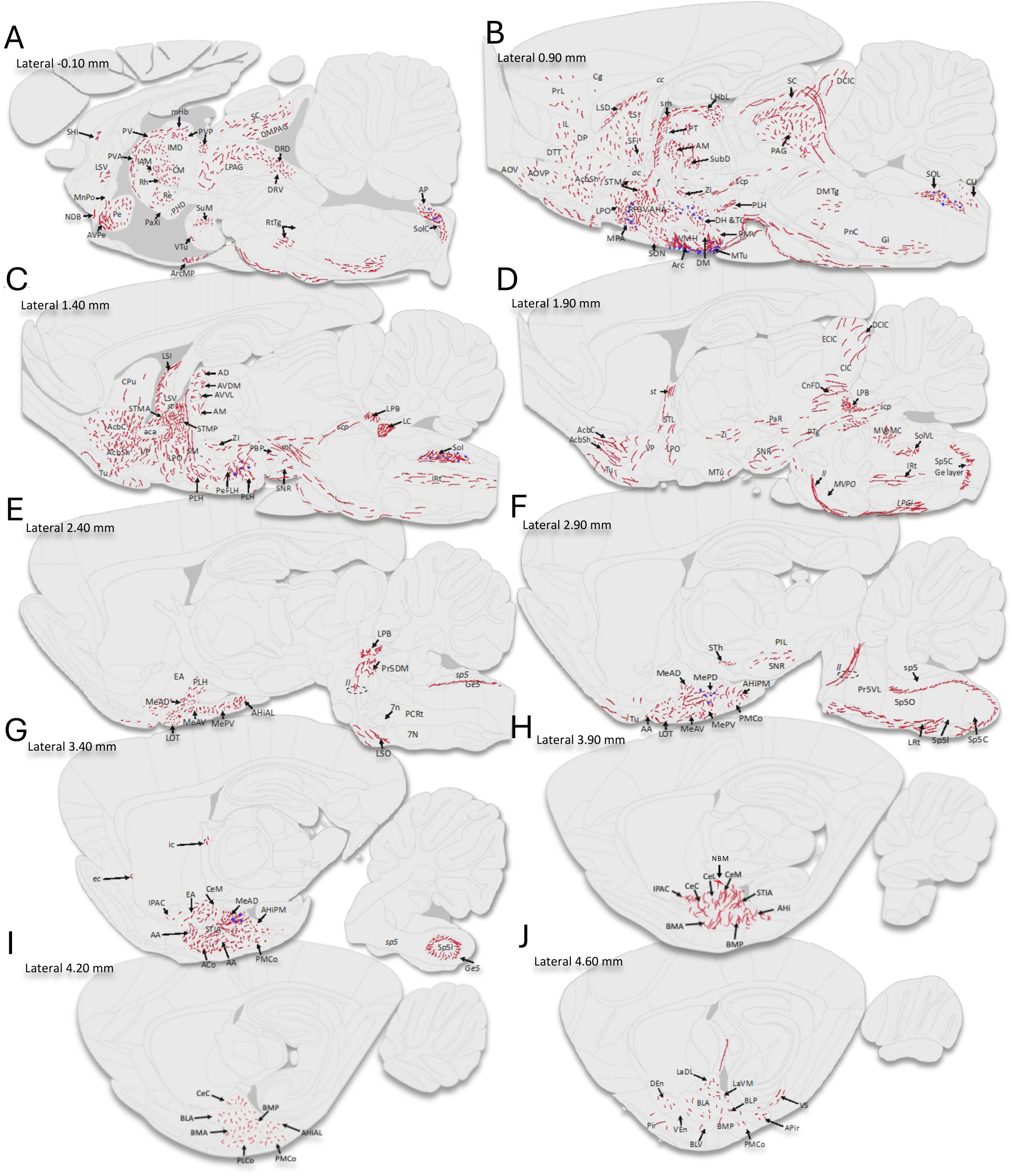
Mapping of KP immunoreactive fibers and cell bodies **in rat brain** in ten septo-temporal planes based on microscopic observations. Blue dots represent KP-ir cell bodies and red lines represent KP-ir fibers. Relevant anatomical regions have supporting photomicrographs form IHC-DAB reactions using the GQ2 anti-kisspeptin antibody. See Table 1 for abbreviations.

In Figure 1, we show KP immunohistochemical reactivity (-ir) in regions, including anteroventral periventricular nucleus (AVPV) and ventromedial preoptic area (VMPO), where KP functions to regulate GnRH secretion, in comparison with other brain regions, with equally intense KP innervation, that are relevant to regulation of dan yang, including median preoptic nucleus (MnPO), septohypothalamic nucleus (SHy), medial septal nucleus (MS, panel A2), lateral septal nucleus (LS) and bed nucleus of stria terminalis (BST, which includes the medioanterior, medioventral and lateroventral divisions, STMA, STMV and STLV, respectively) (panel A). In panel A3, KP-ir cell bodies can be seen in the periventricular region (black arrows). Numerous KP- ir fibers are densely distributed at the ventricular wall with several endings observed at the ventricular lining within the ependymal cell layer (red arrows), suggesting a humoral-secretory function of KP signaling. Panel B depicts a coronal section around Bregma -0.15mm, where four main conducting fiber systems can be clearly seen, i.e. the anterior commissure (ac), the stria medullaris (sm), the stria terminalis (st) and the fornix (f). KP-ir fibers running inside these conducting systems are indicated with red arrows. The dense innervation pattern of KP-ir fibers in the paraventricular anterior nucleus of the thalamus is shown (panel B). In panels C and D we show examples of KP-ir cell bodies and fibers in amygdalar complex. KP-ir fibers can be seen inside the stria terminalis (st), bed nucleus of stria terminalis, intra-amygdalar division (STIA), baso-medial amygdala (BMA), nu. basalis of Meynert (NBM) and medial amygdala (MEA). Interestingly, some fibers within the medial amygdala, postero-dorsal division (MEApd), are observed coming from the stria terminalis, with main axons branching within the MEApd, and possibly originating from the hypothalamus. We observed fibers reaching to amygdala, through the stria terminalis (C1, red arrows). This observation is also supported by the analysis from the Allen Mouse Brain Connectivity Atlas (see Table 1, last two columns entitled "Axons from RP3V viral tracers" and "Axons from Arc viral tracers" and in the SI Figure 2). Panels E and F show the microphotographs taken from the hypothalamic tuberal region, where the KP-ir cell bodies are seen in the posterior division of the arcuate nucleus (ArcP), in premammillary nucleus, ventral division (PMv). KP-ir fibers are observed in other regions relevant for behavioral state control at this level, such as ventral tuberomammillary nu (TMv), magnocelular tuberomammillar nu. and dorsal tubero-mammillary nu., posterior hypothalamus, zona incerta (ZI) and subthalamic nu. (STN) (E). E2 shows the third ventricle at the level of Arc posterior. Abundant KP-ir fibers extended toward the lumen of the ventricle (also see A3, red arrows). Panel G-K are micrographs of samples from parasagittal sections of mouse hindbrain. G shows the KP-ir cell bodies and dense KP-ir fibers in the solitary tract (sol) nu. (NTS) of mouse medulla. H and I show the dense innervation pattern of KP-ir fibers in pontine nuclei: lateral parabrachial nu. (LPBN) and locus coeruleus (LC) respectively. J and K show the dense KP-ir fibers in the spinal trigeminal tract (Sp5, K) and in the gelatinous layer of Sp5 (Ge5). Note that within the Ge5, the KP-ir fiber make perisomatic contacts with the cell bodies of the Ge5 (J, red arrows).

In Figure 2, we present chartings of mouse KP cell and fiber distribution in register with a previous study(14) for purposes of comparison. Panels A, B and C show intense KP innervation of the prefrontal structures, such as cingula cortex (Cg), dorsal peduncular cortex (DP), dorsal tenia tecta (DTT), infralimbic cortex (IL), prelimbic cortex (PrL), ventral orbital cortex (VO), nucleus accumbens shell (AcbSh), medial septal nucleus (MS), nucleus of the diagonal band of Broca (NDB) and dorsal division of the lateral septal nucleus (LSd), which were not previously reported. We also observed abundant innervation in thalamic structures such as nucleus reuniens (Re), periventricular anterior nucleus (PVA) and the epithalamic lateral habenula (LHb). In the amygdaloid complex, we observed KP innervation in cortico-amygdala (CoA), nucleus of lateral olfactory tract (NLOT), medial amygdala (MEA), mainly in the postero-dorsal division, (MEApd), basomedial amygdala (BLA), basolateral amygdala (BLA) and nucleus of the stria terminalis, intra-amygdala division (BSTIA) (Fig. 2, D-G). Midbrain and hindbrain sensorial relay and state control centers, such as premammillary nucleus (PM), dorsal raphe nucleus (DR), periaqueductal gray (PAG), subthalamic nucleus (STh) parabrachial complex (PBc), locus coeruleus (LC), Koelliker-Fuse nucleus (KF) and nucleus of solitary tract (NTS) all possess KP-ir fibers not reported in the previous paper (Fig. 2, H-L).

Kisspeptin (KP) innervation patterns in rat brain are presented in Figure 3. This figure maps KP-immunoreactive fibers (red lines) and cell bodies (blue dots) across ten septo-temporal levels of the rat brain. We observed a pattern of KP fiber distribution similar to that in the mouse brain, particularly in the hypothalamic regions. However, some qualitative differences were noted in the density and extent of KP innervation (see Table 1 for comparison between columns with titles "irGQ2 fiber rat" vs. "irGQ2 fiber mouse". A "++"-“++++” in bold indicates stronger expression in rat than mouse. The differences are especially apparent in the telencephalic and diencephalic regions, where a generally stronger KP expression was observed in rat brain. These regions include the cingulate cortex, dorsal peduncular cortex, dorsal tenia tecta, infralimbic cortex, prelimbic cortex, ventral orbital cortex, olfactory area, anterior olfactory nucleus, olfactory tubercle, extended amygdala, anterior amygdala, and accumbens shell. These differences emphasize the importance of comparative studies to understand the conserved and divergent roles of KP signaling across different species, particularly in non-reproductive functions, where KP signaling may have undergone exaptation processes allowing it to serve other relevant functions for the species in question (40).

### 3.2 Detailed description of regional distribution of KP fibers with an emphasis on sensorial processing and behavioral state control

The distribution of kisspeptin fibers in several regions involving numerous crucial central sensorial relays and behavioral-state control structures(20–25) suggests wider participation of this peptide in other functions besides its known participation in reproduction and energy balance. In the following sections, we describe these regions in detail.

#### 3.2.1 Telencephalon

In rodent telencephalon, the most prominent KP fiber-expressing regions are located in the *lateral septal region* in male and female mice and rats. The strongest expression was observed in the dorsal lateral septal nucleus (LSD) (Fig. 1A & A2, Fig. 2B-D, and Fig 3B-C), followed by ventral LS (LSV) (Fig. 2C, and 3C). LS is involved in the integration of sensorial and emotional information, and can modulate the activity of other brain regions involved in the regulation of behavioral states, such as the hypothalamus, amygdala and hippocampus(41, 42). Recent studies have reported the main afferents of the above regions are the subdivisions of the hippocampal formation and the main efferent regions of the LSD are the ventral tegmental area and the supramammilary nucleus, whereas the LSV projects mainly to the medial hypothalamic area and medial preoptic area(42). Activity of lateral septum can lead to changes in behavioral state, including increased exploratory behavioral, social interactions and decreased anxiety, particularly in response to social and emotional stimuli. Such behavioral changes have been reported in clinical studies with KP infusion in human subjects(43). LS is also a key player for arousal and wakefulness, as well as for regulating processes related to mood and motivation through its extensive connections with numerous regions such as the hippocampus, amygdala, hypothalamus and with the mesocorticolimbic dopamine system(44).

The *medial septum* (MS) also contained scattered and sparse innervation in rats and mice (Fig. 1 A and 2 B). This structure is crucially involved in regulating hippocampal oscillations(45, 46), i.e. the rhythmic pattern of neuronal activity detected via changes in local field potential, which have been found to be relevant for the encoding of information and supporting processes such as learning and memory(47). Recent studies have shown that the medial septum acts as a neural hub that regulates theta oscillations and locomotion speed, influencing goal-directed behaviors such as spatial navigation (48). Other behaviors influenced by the medial septum include sleep, attention, learning and memory, motivation and emotional processing(49–51).

Sparse fibers were found in septofimbrial nucleus (Fig. 3B), which is an important source of cholinergic inputs to the hippocampus and the entorhinal cortex, that influence hippocampal plasticity and learning and memory under emotional stimuli; and the septohippocampal (SHi, Fig. 2B) nuclei that provide important GABAergic inputs to hippocampus playing a role in the generation of hippocampal theta oscillation(52). The above two KPergic regions are described for the first time in this study.

Similar to Brailoiu et. al(39), we found sparse fibers in the horizontal and vertical limbs of the nucleus of diagonal band of Broca (NDB) (Fig. 2B). The diagonal band of Broca interconnects the amygdala and the septal area and projects particularly to the CA2 subfield of hippocampus, which is involved in memory retrieval and social functions (53, 54). It contains the second largest cholinergic neuronal population in the brain (after the nucleus basalis of Meynert)(55). It is particularly important in the aging of the brain, since the cholinergic neuronal loss in dementias has been commonly observed in human, suggesting a possible connection between KP neuromodulation and neurodegeneration-associated cognitive dysfunction.

An interesting finding of this study was the remarkable presence of kisspeptin fibers in the *nucleus accumbens,* in both core and shell divisions (AcbC and AcbSh), with a low density in mice and a moderate density in rats (Table 1, Fig. 2A and Fig. 3B-D). AcbC receives inputs from the prefrontal cortex, the amygdala, and the ventral tegmental area (VTA), which is a key component of the brain’s reward system. The AcbC integrates these inputs to evaluate the salience and value of rewards and helps to regulate motivated behavior accordingly. The nucleus accumbens shell is an important brain region involved in the processing of reward-related information and the regulation of motivated behavior, including food intake, drug seeking, and social behavior(56).

Sparse fibers were observed in the caudate putamen (CPu Fig. 2D & 3C), ventral pallidus (VP, Fig. 3D), the sustantia innominata (SI), and in main fiber pathways such as the internal capsule (ic) and external capsule (ec).

In telencephalic neocortical regions, we found sparse fibers distributed through the dorsal tenia tecta (DTT) and the dorsal peduncular (DP), prelimbic (PrL) , infralimbic (IL), and cingular (Cg) cortices, regions not reported in previous studies. The observed fibers were mainly located in layers 2/3, and no differences between species were observed (Tab. 1 and Fig. 2A & Fig. 3B). The prelimbic (PrL), infralimbic (IL) and cingulate cortices (Cg) have been classically considered to be involved in integrating the role of context in the expression of fear- or reward-induced behaviors(57–59). The DTT and the DP have been classically involved in olfactory information processing, and recently these two structures have been shown to send inputs with information related to psychosocial stress to the dorsomedial hypothalamus, where a cardiovascular and thermogenic sympathetic stress response can be integrated(60). In other areas classically related to the processing of olfactory information, such as the ventral and ventroposterior of the anterior olfactory nucleus, (AOV and AOVP) and the olfactory tubercle (Tu), we found sparse fibers only in the rat (Tab. 1, Fig. 3B-D).

In the amygdalar complex, we confirmed the KP-ir cell population mainly confined within the medial amygdala’s postero-dorsal division (MePD) which has been reported already elsewhere (61, 62) (Table 1 & Figs. 1C-D, 2F-G and 3F-G). ISH study in gonadally intact rats and mice showed that *Kiss1* levels were higher in males than females and fluctuated with the estrous cycle, peaking at proestrus in rats (63) suggesting its potential influence reproductive and broader brain functions. Kisspeptin- immunolabelled neurons in the MePD and an axon terminal process were observed to innervate the adjacent nucleus of the stria terminalis intra-amygdaloid division and the stria terminalis. Also, we observed KP-expressing cell bodies in the septo-hippocampal (SHi) and septo-fimbrial (SFi) region surrounding the anterior commissure, and extending into the septum. These neurons were characterized neurochemically in mice in our previous report(1).

In relation to density of fibers in the septo-bed nucleus of stria terminalis complex, regions with a moderate/intense density of fibers were found within the septal part of the bed nucleus of the stria terminalis (BSTMA, BSTMP, BSTLV and BSTLP, Fig. 1A; Fig. 2B-D and Fig. 3B-C). The temporal intra-amygdala division of the BST (BSTIA, Fig. 1C; Fig. 2F and Fig. 3H) showed a selective expression of KP fibers. Other regions in the extended amygdala that also showed low/scattered expression of KP fibers were the central amygdalar nucleus (CeC and CeM Fig. 3G-H) and the medial amygdalar nucleus (MePD and MePV Fig. 2,F-G and Fig. 3.G-H). Sparse fibers were found in the oval nucleus of the stria terminalis (BSTLD), the basolateral amygdala (BLA), the basomedial amygdala (BMA), the lateral part of the central amygdalar nucleus (CeL), the cortico-amygdala (CoA) the nucleus of the lateral olfactory tract (NLOT) and the stria terminalis (st, Fig. 1C). Compared with the study of Brailou et al, here we report the presence of fibers in the anterior, medial, and cortical amygdala and the intra-amygdaloid division of the stria terminalis (Tab. 2).

**Table 2.** Semiquantitative analysis of KP immunopositive cells and fiber density in regions hosting KPergic neurons and regions related to sensorial-brain state processing in old female and male mice

|  | FC | OVX | FC vs OVX | MC | ORX | MC vs ORX | FC vs MC |
| --- | --- | --- | --- | --- | --- | --- | --- |
| AcbSh | +++ | +++ | FC ≈ OVX | ++ | ++ | MC ≈ ORX | FC > MC |
| Arc | +++ | +++ | FC ≈ OVX | ++ | ++ | MC ≈ ORX | FC > MC |
| BSTam | +++ | +++ | FC ≈ OVX | ++ | ++ | MC ≈ ORX | FC > MC |
| DB | +++ | +++ | FC ≈ OVX | + | ++ | MC ≈ ORX | FC > MC |
| DH | + | + | FC ≈ OVX | ++ | ++ | MC < ORX | **FC < MC** |
| LC | ++++ | ++++ | FC ≈ OVX | ++++ | ++++ | MC ≈ ORX | FC ≈ MC |
| LSD | + | + | FC ≈ OVX | + | + | MC ≈ ORX | FC ≈ MC |
| MEA | + | + | FC ≈ OVX | + | + | MC ≈ ORX | FC ≈ MC |
| MS | ++ | ++ | FC ≈ OVX | + | + | MC ≈ ORX | FC > MC |
| NTS | ++++ | ++++ | FC ≈ OVX | ++++ | ++++ | MC ≈ ORX | FC ≈ MC |
| PaAP | +++ | +++ | FC ≈ OVX | ++ | ++ | MC ≈ ORX | FC > MC |
| PBN | ++++ | ++++ | FC ≈ OVX | ++++ | ++++ | MC ≈ ORX | FC ≈ MC |
| TM | + | + | FC ≈ OVX | ++ | ++ | MC < ORX | **FC < MC** |
| PMv | + | + | FC ≈ OVX | ++ | ++ | MC ≈ ORX | FC < MC |
| PVA | ++ | ++ | FC ≈ OVX | +++ | ++++ | MC < ORX | FC < MC |
| PVN | +++ | +++ | FC ≈ OVX | ++ | ++ | MC ≈ ORX | FC > MC |
| RP3V | +++ | ++++ | FC < OVX | ++ | ++ | MC ≈ ORX | FC > MC |
| AH (adj SCN) | +++ | +++ | FC ≈ OVX | ++ | ++ | MC ≈ ORX | FC > MC |
| SHy | +++ | ++++ | FC < OVX | ++ | ++ | MC ≈ ORX | FC > MC |
| STIA | + | + | FC ≈ OVX | + | + | MC ≈ ORX | FC ≈ MC |
| Wall 3V | +++ | +++ | FC ≈ OVX | + | + | MC ≈ ORX | FC ≈ MC |
| ROIs adj.<br>(f/st/sm/ac) | ++ | ++ | FC ≈ OVX | ++ | ++ | MC ≈ ORX | FC > MC |

#### 3.2.2. Diencephalon

The *hypothalamus* contains four of the seven KP neuronal populations described in detail here and in a companion report(1), i.e. the rostral periventricular (RP3V) population mainly located in the medial preoptic area (MPO) and in the anteroventral periventricular area (AVPe); the arcuate (Arc) population; the dorsomedial hypothalamus (DMH); and the ventral premammillary nu. (PMv) (Fig. 1A; Fig. 2 C, D, F, G, H and Fig. 3B). After a careful analysis of immunoreacted serial sections, and following the pathways of the labelled fibers, we identified some extrahypothalamic regions likely to be targets of these regions. Ascending fibers originating in the RP3V population travel lateral-dorsal-caudally toward the stria medularis (sm), and the stria terminalis (st) conduction systems (Fig. 1, B). In their path, they extensively innervate the bed nucleus of the stria terminalis (Tab. 2). It is worth noting the presence of KP fibers innervating the border between the *medial (MHb) and lateral (LHb) habenula* (Fig. 2, F-G and Fig. 3 B), as it is known that the main input to the habenula is the stria medullaris (*sm*). Other fibers originating in the MPO/AVPV region travel to innervate the organum vasculosum of the lamina terminalis (OVLT) and then ascend dorsally and laterally to innervate the *accumbens core* (AcbC) and the *lateral septum* regions.

We observed high-moderate densities of KP-ir fibers (see Tab. 2) in the caudal periventricular area and ventral periventricular areas of the arcuate nucleus (Arc, Fig. 1E, and Fig. 2F-H). Some targets of the KP neural population in the posterior hypothalamus were identified by following the labelled axon in serially collected sections in coronal and parasagittal planes Some fibers originating in the Arc travelled in the base of the brain reaching the internal capsule (ic) and optic tract through which they travel in a path to the subthalamic nucleus (STh), the lateral part of the supramammillary nucleus (SuML) and continuing to the central nucleus of the amygdala (CeA). Some efferent fibers traveled caudally to innervate the ventral premammillary nucleus (VPM), the ventral tuberomammillary nucleus (VTu), and the supramammillary nucleus (SuM). Additional fibers surrounded the ventromedial hypothalamus (VMH) and travelled dorsally to innervate the anterior hypothalamic area, continuing on to innervate medial nuclei of the thalamus including the paraventricular thalamic (PVT). The latter showed the highest innervation, albeit the intermediodorsal (IMD) the centromedial (CM), mediodorsal (MD), paratenial (PT), rheuniens (Re) and rhomboid (Rh) nuclei of the thalamus also contained KP-ir fibers.

The population of weakly KP-immunopositive cells scattered in the medio-lateral dorsal hypothalamic area (DH) spanning from the posterior part of the anterior hypothalamus (AHp) to the posterior hypothalamus (PH), and centered in the tuber cinereum area (TC) (Fig. 1F; Fig. 2F-H, and Fig. 3B) were observed to project to adjacent zona incerta (ZI). Finally, the last population of KP neurons in the PMv was very weakly labelled and it was not possible to identify the axonal projections.

Other regions in the hypothalamus containing KP fibers are the median preoptic nucleus (MPO), the parastriatal nucleus (PS), the paraventricular nucleus medial magnocellular division (PVNmm) and the parvocellular division (PVNmp), and in rat also the ventral premammillary nucleus (PMv), ventral tuberomammillary nucleus (VTu), and suprammamilary nucleus (SuM). A low density of fibers was observed in dorsomedial hypothalamic nucleus, MPO, lateral preoptic area, periventricular hypothalamic nucleus anterior (PVa), and supraoptic nucleus (SO). Sparse fibers were found in the dorsal hypothalamus (DH), zona incerta (ZI), lateral hypothalamic area (LH), posterior hypothalamic area (PH), retrochiasmatic area (RCh), and ventromedial hypothalamic nucleus (VMH). These observations confirm and extend extensive investigations of the hypothalamic KP system(16).

In the thalamus, the highest density of fibers was observed in the paraventricular thalamic nucleus (PVT) in rats and mice (Fig. 1B; Fig. 2E-F and 3B). Low fibers densities were also observed in other nuclei of the thalamus, such as in the centromedial (CM), mediodorsal (MD), paratenial (PT), reuniens (Re), and rhomboid (Rh). The epithalamic habenular nucleus (Hb) showed a moderate density of fibers mainly located in the border between its medial and lateral subdivisions (Fig. 2F-G and 3B ), with some of these fibers arriving through the stria medullaris.

#### 3.2.3. Mesencephalon

Within the mesencephalon, Brailoiu et al(39) reported a sparse number of kisspeptin-immunoreactive fibers only in the dorsomedial periaqueductal gray substance (DMPAG). In our study this region had the highest density of kisspeptin fibers within the mesencephalon (Fig. 2J and Fig. 3B). However, we also visualized scattered fibers in the central nucleus of the inferior colliculus (CIC, Fig. 2I and Fig. 3D), dorsal raphe nucleus (DR, Fig. 2J and Fig. 3A), and sparse fibers in the deep mesencephalic nucleus (DpMe), interpeduncular nucleus (IP), median raphe nucleus (MnR), substantia nigra (SN), intermediate and deep gray layer of the superior colliculus (DpG), supramammillary nucleus (Sum), substantia nigra pars reticulata (SNpr) and ventral tegmental area (VTA). The functional implications for those Kp projections are not yet ascertained.

#### 3.2.4. Metencephalon and myelencephalon

In the metencephalon and myelencephalon, we observed kisspeptin immunopositive cell bodies in the *nucleus tractus solitarius* (NTS or Sol), and *area postrema* (AP) Fig. 1 G; Fig. 2L and Fig. 3A-D). We did not observe immunopositive neurons in the caudoventrolateral or the lateral reticular nuclei (CVL and LRt), nor the spinal trigeminal tract, as reported in Brailoiu et al(39). With respect to the density of the fibers, intense labelling was found in the lateral parabrachial nucleus (LPB, Fig. 1H, Fig. 2K), the Kolliker-Fuse nucleus (KF Fig. 2K), NTS, (Fig. 1G; Fig. 2L and Fig. 3A-D) in the dorsomedial part of the principal sensorial component of the trigeminal nucleus (Pr5DM, Fig. 1K and Fig. 3D-G) and locus coeruleus (LC, Fig. 1I; Fig. 2K and Fig. 3C). The rat brain also showed intense Kp labeling in the gelatinous layer and the spinal nerve 5 (Fig. 1J-K and Fig. 3E-F). Moderate density was observed in the superior cerebral peduncle (scp) . Low intensity was observed in several nuclei of the reticular formation, including the caudoventral reticular nucleus (CVL), the gigantocellular reticular nucleus (Gi), the intermediate reticular nucleus, the lateral reticular nucleus (LRt), the medullary dorsal reticular nucleus (MdD), the parvicellular reticular nucleus (PCRt), and the rostroventrolateral reticular nucleus (RVL). Sparse fiber density was observed in the ambiguus nucleus (Amb); lateral superior olive (LSO); ventral medullary reticular nucleus (MdV); in subregions of the pontine nucleus such as the brachium pontis (bpPn), the reticular oral part (PnO), and the reticular caudal part (PnC); and in the raphe magnus (RMg), obscurus (Rob), and pallidus (RPa).

### 3.3 KP fiber density: sex differences and short- and long-term effects of gonadectomy

Numerous studies have reported that gonadal status affects *Kiss1* expression, with KP^Arc^ expression decreased, and rostral hypothalamus KP^RP3V^ expression increased by gonadal steroids(64–66). We have also reported male and female *Kiss1* expression differences and acute sensitivity to GNX (i.e. within 4-7 weeks of GNX)(1).

We extensively studied the fiber distribution in male and female mice that underwent gonadectomy in young adulthood (3 month old). The experimental subjects were perfused/fixed at ages of 4-5 months (1-2 months after GNX) or 15 months (12 months after GNX). We examined KP-ir fibers densities aiming to assess if there are sex differences in KP fiber distribution and whether castration modifies the density of fibers apparent both shortly after GNX, and in aged mice after early GNX, compared with intact subjects of the same age.

Supplementary Information Figure 1 shows the cumulative (summed) fiber density charting of three young adult animals from each of four groups: male/female intact and male/female gonadectomized. The chartings were made under microscopical observation and verified by two junior and two senior experimenters of this research team. The cumulative fiber densities displayed in the SI Figure 1 show no appreciable differences among the four groups following careful analysis of KP-ir fiber density across groups. This may be due to a lack of regulation of KP production in any of the neuronal groups contributing nerve terminals to the assessed areas, or, possibly, compensatory and opposite regulation of KP production among multiple contributing KPergic inputs to a given region.

We further made an analysis of KP Cre-driver in ARC and RP3V tracing experiments performed by the Allen Brain Institute and deposited in the Allen Brain Atlas (showed in the last two columns of the table 1). In the supplementary information (SI) figure 1, we highlight the projection targets of R3PV or Arc *Kiss1*-Cre neurons transfected with AAV. The data from two *Kiss1*-Cre mice injected in the arcuate nucleus (Arc, experiments 232311236 and 232310521) and two *Kiss1*-Cre mice injected in the rostral periventricular region (R3PV, experiments 299247009 and 301989585) are reproduced in this Figure. The VGAT-expressing *Kiss1* R3PV neuronal population and its projections are represented in red, and the VGLUT2-expressing *Kiss1* Arc population and its projections are represented in green. The different views show a reciprocal dense innervation between the Arc and the RP3V. Additionally it is clear that the majority of regions in the striatum, septum, BNST, thalamus, hypothalamus, and amygdala receive convergent innervation from both Arc and RP3V Kiss1 neuronal populations.

According to our KP fiber study, and analysis of the Allen Connectivity Map, a representation of the relative fiber density distribution and some identified kisspeptin pathways through the mouse brain is presented in the flat map in SI Figure 3. Most of the regions involved in sensorial processing, motor integration and behavioral state control that were described and discussed previously are shown in this representation with reciprocal or convergent innervations from both KP-RP3P and KP-Arc.

Table 2 is a report of a semiquantitative analysis of 22 regions from 12 15-month old mice, within the scope of this study. In female subjects, 19 of the 22 regions were found similar in term of density of KP-ir fibers between old intact ("FC" stands for female control) and experimental ("OVX" stands for ovariectomy performed 12 months prior to perfusion/fixation) female mice. The septohypothalamic nucleus showed increased KP-ir fiber density in OVX subjects. This region, beside hosting *Kiss1/Slc32a1/Ar* expressing neurons(1) (Fig. 1, A and A1), receives direct RP3V^KP^ and Arc^KP^ projections as shown by Allen Mouse Brain Atlas (SI figure 1). Hence, the source of the neuronal cell group contributing to the increased KP-ir fiber density in septohypothalamic nucleus after OVX is not clear. In the RP3V and MPO region, the samples from OVX subjects also showed an increased KP immunoreactivity (Fig. 4). This region, besides hosting a KP neuronal population itself, also receives Arc^KP^ axonal projections, as demonstrated in SI Figure 1. In male subjects, 18 of the 22 regions showed no apparent differences between intact ("MC" stands for male control) and orchiectomized ("ORX" stands for orchiectomy) male mice. Three regions exhibited higher densities of KP-ir fibers in the ORX group compared to the aged-matched intact controls: dorsal hypothalamus (DH, Fig. 4), paraventricular anterior thalamus (PVA) and tuberomammillary nucleus (TM). Also, in these three regions, the MC group showed higher densities than the FC group (Fig. 4).

**Fig. 4.**
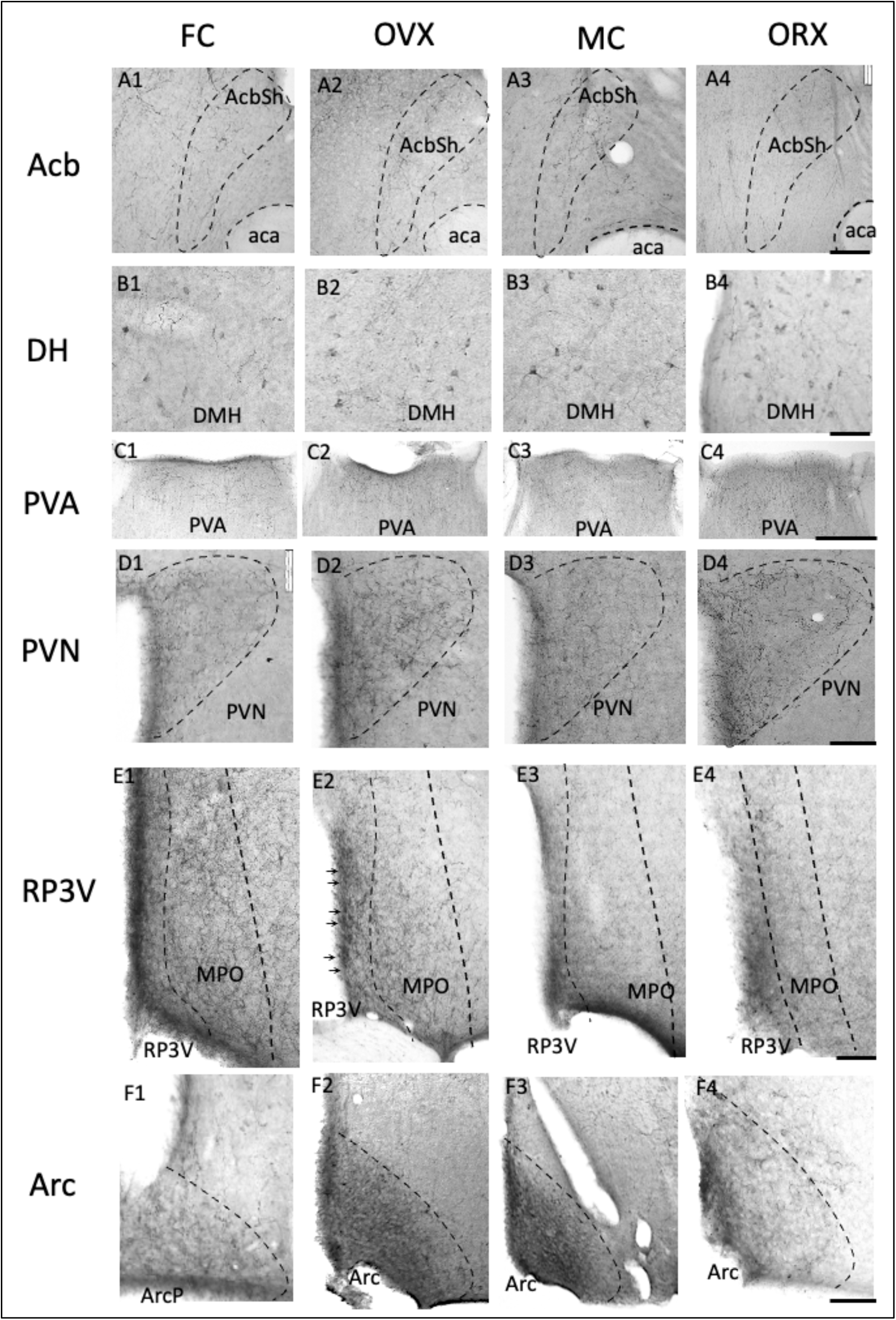
Examples of KP-ir fiber density in old mice: comparison of four groups. FC: samples from female control, intact mice of 15-month-old; OVX: female mice undergoing ovariectomy in young adulthood (3- month-old) and perfused/fixed 12 months after OVX); MC: male intact mice; ORX: male mice undergoing orchiectomy in young adulthood (3 month old) and perfused/fixed 12 months after ORX). Acb, nucleus accumbens; AcbSh, nucleus accumbens shell: Arc, arcuate hypothalamic n.; ArcP, arcuate hypothalamic n. posterior; DH, dorsal hypothalamus; DMH, dorsomedial hypothalamus; PVA, paraventricular nucleus anterior division of the thalamus; PVN, paraventricular nucleus of the hypothalamus; RP3V, rostro- periventricular complex of third ventricle of the hypothalamus. Scale bars: 100µm.

### 3.4 Mapping of *Kiss1r and its co-expression with Slc32a1, Slc17a6 and Kiss1* with RNAscope methods: assessment of *Kiss1r* distribution in CNS and assessment of Kiss1-Kiss1r co-expression

Using RNAscope dual and multiplex ISH methods, we report here (Fig. 5) semi-quantitative expression levels of *Kiss1r*, depicted as different intensities of blue shading, in mouse brain sagittal serial sections. In contrast to the discrete expression of *Kiss1* in seven neuronal populations in brain(1), *Kiss1r* expression is diffuse and widespread. *Kiss1r* was preferentially expressed in non-GABAergic neurons (Table 3), such as hippocampal projection neurons, mainly glutamatergic thalamic nuclei, mammillary complex neurons, cerebellar granule cells, and the interposed nucleus. *Kiss1r* was also co-expressed with *Kiss1* itself in hypothalamic neurons, suggesting an autocrine component of KP signaling in the hypothalamus. For example (Fig. 6, panel 14 and 17) *Kiss1r* and *Kiss1* are co-expressed in RP3V, using 4-channel RNAscope LS Multiplex Fluorescent Assay against mRNA for kisspeptin receptor (*Kiss1r*, red dots), kisspeptin (*Kiss1*, green dots, VGAT (*Slc32a1*, blue dots) and VGLUT2 (*Slc17a6*). In RP3V (Fig. 6, panel 14), six out of nine cells (6/9) in a representative confocal microphotograph of this area co-expressed the four mRNAs (indicated with quadri-color-arrows). This is in accordance with our previous observation (1) that the kisspeptinergic neuronal population in the RP3V co-express VGAT and VGLUT2 mRNA, indicating the GABA/Glutamate co-transmission nature of those subcortical neurons. Three cells not expressing Kiss1+ but expressing Kiss1r were Slc32a1^-^ and Slc17a6^+^. In Arc (Fig. 6, panel 16), Kiss1r was also found co-expressed in Kiss1- and Slc17a6-expressing cells lacking expression of Slc32a1 (indicated with tricolor arrows in the panel 16). Insets show 4-channel separated screenshots of the cell in the center. The red thin arrows, generated by the Leica Application Suite (LAS) software, indicate the very same coordinates in each channel. While previous studies(31, 67) have reported exclusive expression of Kiss1 and Kiss1r in hypothalamus, careful inspection of the reported data suggest that co-expressing cells may be undercounted without optimal *colocalization at the cellular level* and employment of precise and enhanced image processing tools. Panel 12 of Fig. 6, depicts expression of Kiss1r in each of the cells in the ventral division of the premammillary nucleus (PMv), a region recently reported to host the newly discovered KP population. Most neurons neighboring *Kiss1-expressing* cells also expressed *Kiss1r*, suggesting that the KP/Kiss1R pathway may use *paracrine* mechanisms, as well as autocrine ones, in addition to classical neurotransmission through axon innervation and transmitter co-release, and potentially Kp humoral function via release into cerebrospinal fluid (Fig. 1, Panels A3 and E2).

**Fig. 5.**
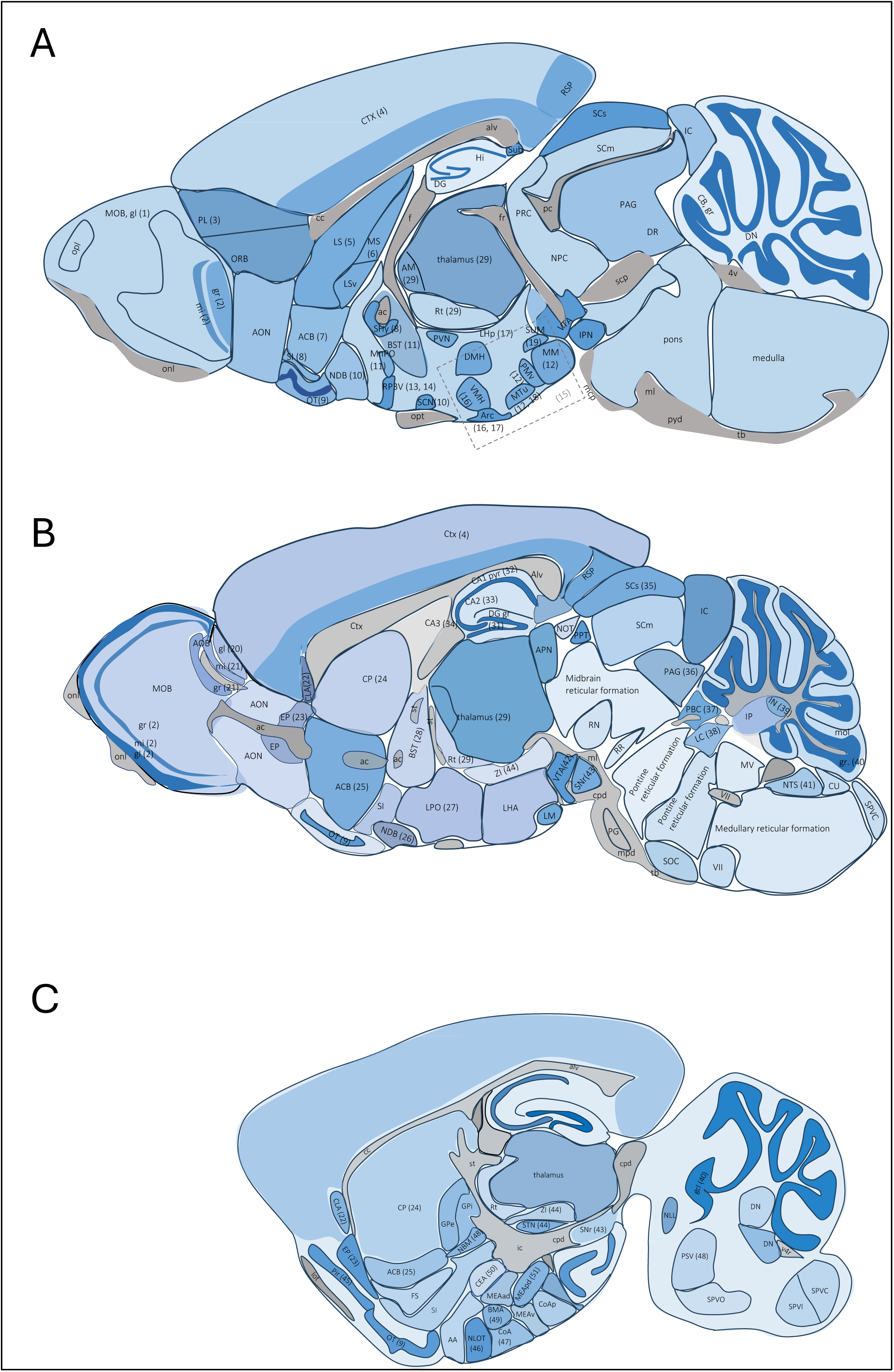
Mapping of Kiss1r (RNA encoding KP receptor) expression. (symbolized by the intensity of blue shading) in three septo-temporal planes (A-C) based on microscopical observation reported in the table 2. Numbers following region-abbreviations correspond to the panel numbers of Figure 6. ACB, nucleus accumbens; AM, anteromedial nucleus of thalamus; AOB gr, granule cell layer AOB; AOB ipl, inner plexiform layer AOB; AOB mi, mitral cell layer AOB; AOB opl, outer plexiform layer AOB; AOB, glomerular layer AOB; MOB, main olfactory bulb; Arc, arcuate hypothalamic n.; BMA, basomedial amygdala; BST, bed nucleus of stria terminalis; CEA, central amygdala; CLA, Claustrum; CoA, corticoamygdala; CP, caudoputamen; DC, dentate gyrus; EP, endopiriform n.; gcl, granule cell layer of DG; gr, cerebellar granule cell layer; LC, locus coeruleus; LPO, lateral preoptic area; LS, lateral septal n. ; MEA, medial amygdala; MnPO, median preoptic n.; MOB, main olfactory bulb; MOB gl, glomerular layer MOB; MOB gr, granule cell layer MOB; MOB ipl, inner plexiform layer MOB; MOB mi, mitral cell layer MOB; MOB opl, outer plexiform layer MOB; Mol, cerebellar molecular cell layer; MS, medial septal n.; NBM, nucleus basalis of Meynert; NDB, diagonal band nucleus; NLOT, nucleus of lateral olfactory tract; NTS, nucleus tractus solitarii; OT, olfactory tubercle; PAG, periaqueductal grey; PB, parabrachial n.; PC, Purkinje cell; PfC, prefrontal cortex; Pir, piriform cortex; PL, prelimbic area; plc, pyramidal cell layer; PMv, premammillary n., ventral; RP3V, rostro-periventricular n. of 3rd ventricle; Rt, reticular thalamic nucleus; SC, superior colliculus; SCN, suprachiasmatic n.; scp, superior cerebellar peduncle; SHy, septo-hypothalamic n.; SNr, substantia nigra reticulata; SS, somatosensorial cortex; STN, subthalamic nucleus; SUM, supramammillary n.; TMv, tuberomammillary n., ventral; VMN, ventromedial hypothalamic n.; VTA, ventral tegmental area; ZI, zona incerta.

**Fig. 6.**
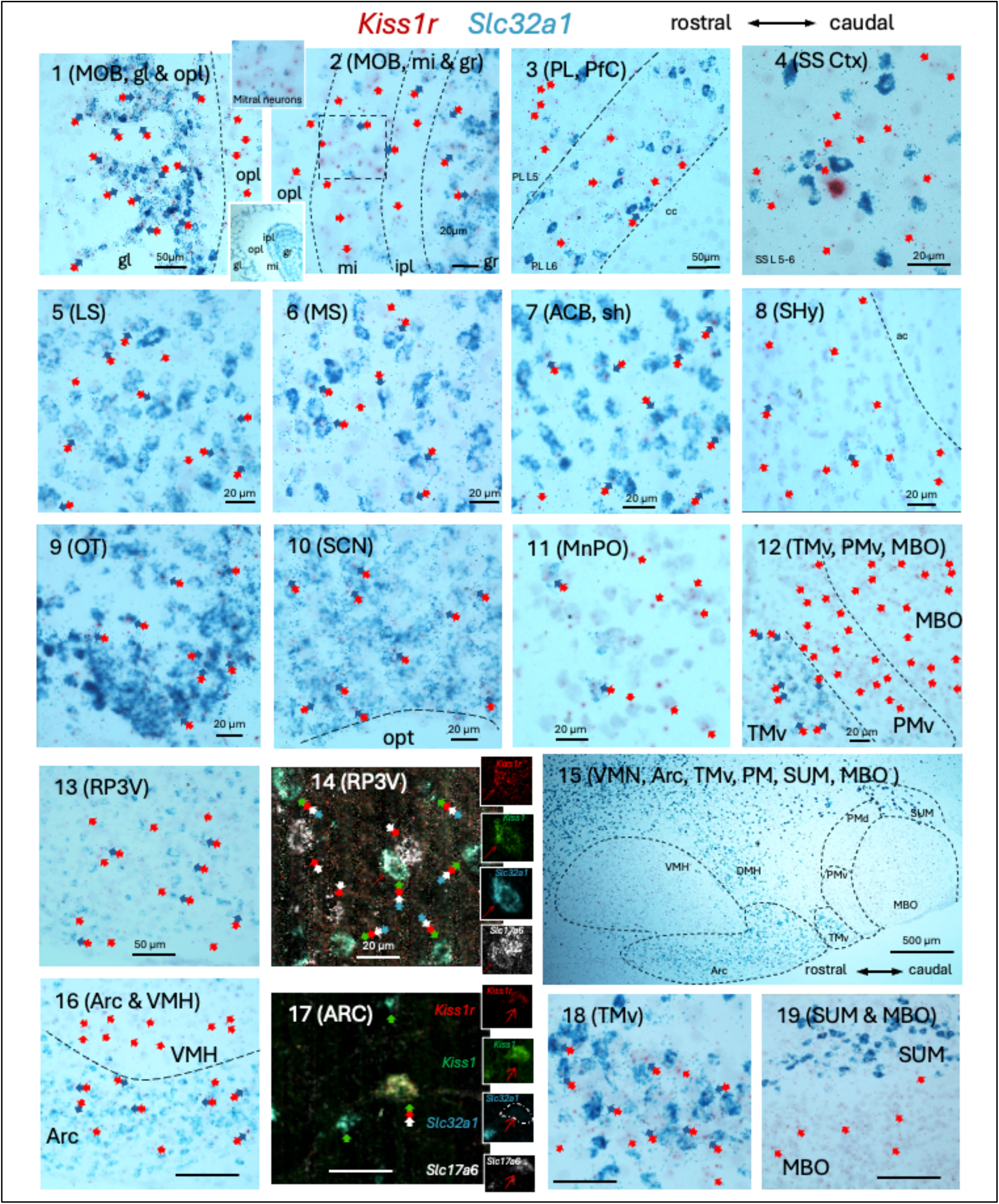
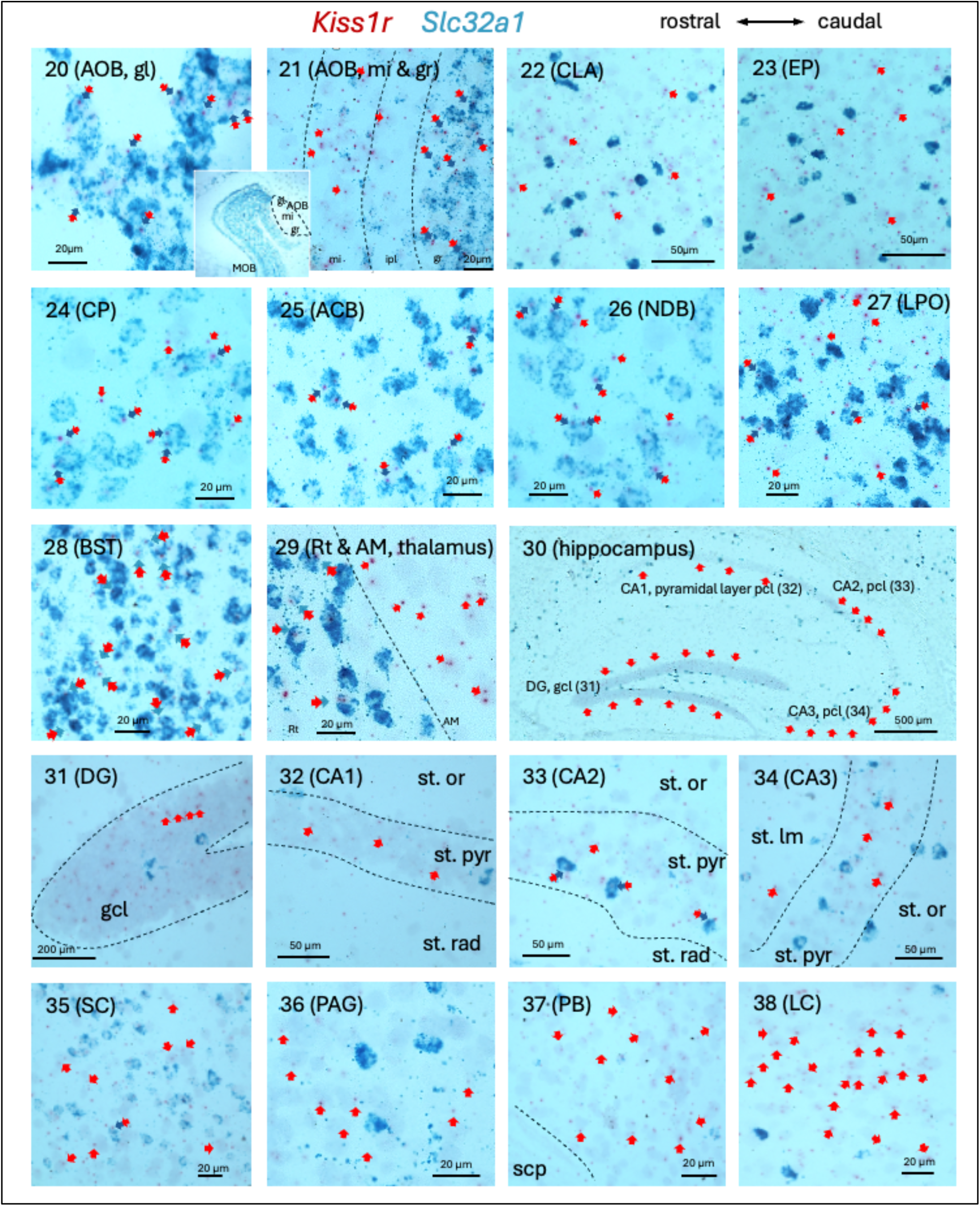
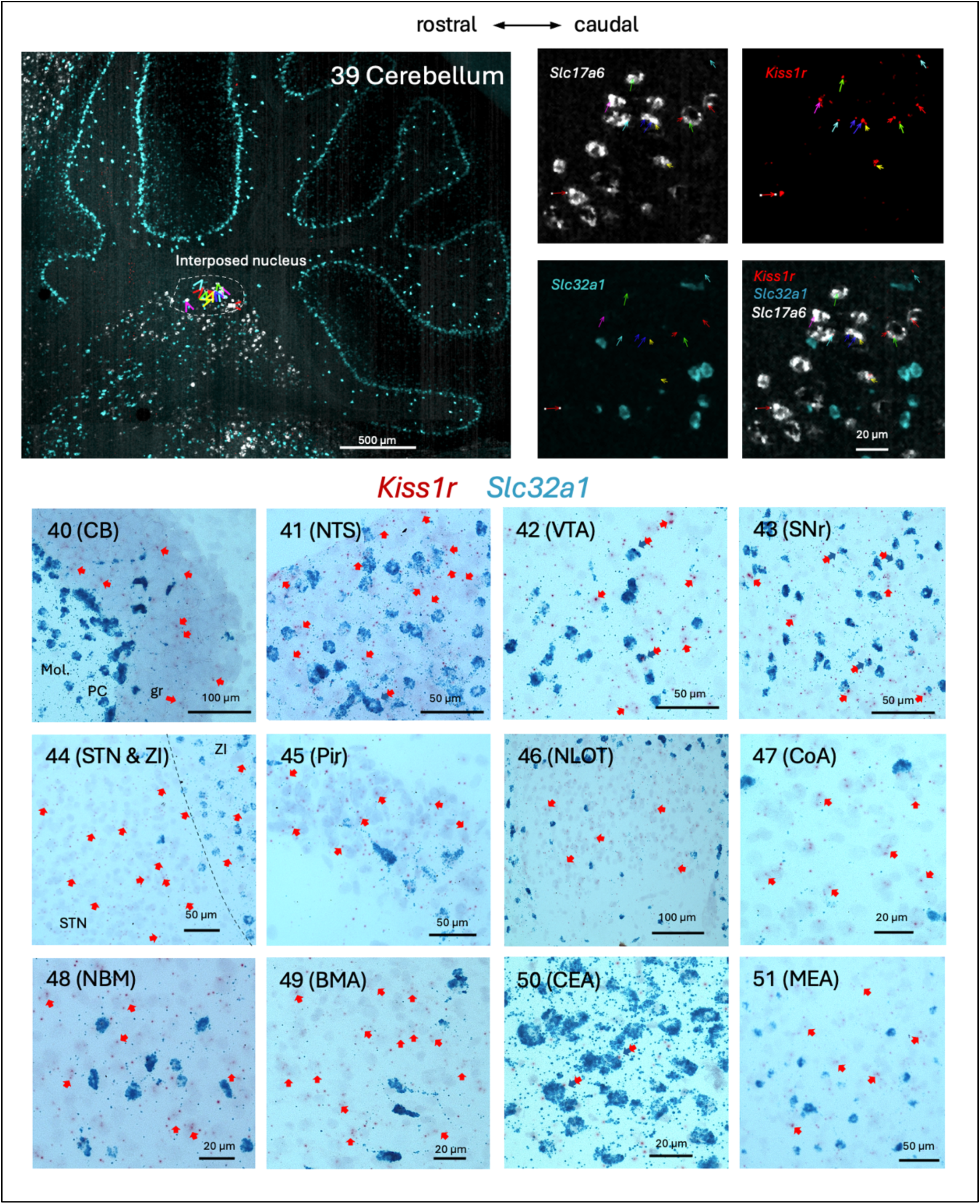
**Examples illustrating Kiss1r co-expression with GABAergic neurons (Slc32a1 mRNA encoding VGAT) in cortical and subcortical regions from sagittal sections**. **Co-expression Kiss1r with Kiss1 (mRNA encoding kisspeptin), Slc17a6 (mRNA encoding VGluT2) and Sla32a1, was examined at the RP3V and Arc regions (panels 14,17, and 39)**. Panels **1** and **2**: MOB, main olfactory bulb; gl, glomerular layer; opl, outer plexiform layer; mi, mitral cell layer; ipl, inner plexiform layer; gr, granule cell layer. N.B. Kiss1r (red dots) is widely expressed in the MOB periglomerular cells which are inhibitory interneurons (Slc32a1, blue patches, co-expressing, bicolor arrows in panel 1). Kiss1r is also widely expressed in the opl, not Slc32a1 expressing cells (red arrows) (panel 1). The principal projection cells in MOB, the mitral cells, mi, are strongly expressing the Kiss1r (panel 2), as well as the GABAergic neurons in the granule cell layer (gr). Panel **3**: PL, prelimbic area of prefrontal cortex (PfC), most Kiss1r labeling (red arrows) are not on Slc32a1 expressing neurons. Panel **4**: somatosensorial cortex (SS Ctx), layer 5-6, most Kiss1r labeling (red arrows) are not on Slc32a1 expressing neurons. In lateral, medial septal nuclei (LS, panel **5**, and MS, panel **6**) and nucleus accumbens (ACB, panel **7**), the Kiss1r was mainly found in Slc32a1 expressing (GABAergic) neurons (bicolor double arrows). Panel **8**: Shy, septo- hypothalamic nucleus, the Kiss1r labeling are main in none-Scl32a1 cells (red arrows). Panel **9** and **10**: OT, olfactory tubercle and SCN, suprachiasmatic nucleus, the Kiss1r is almost exclusively expressed in the Scl32a1 expressing neurons (bicolor double arrows). Panel **11**: MnPO, median preoptic nucleus, Kiss1r is expressed in both GABAergic and no-GABAergic cells. In panel **12**, three adjacent hypothalamic regions are displayed. TMv: tuberomammillary nucleus ventral division, most GABAergic neurons expressed Kiss1; PMv, premammillary nucleus, ventral division, and MBO, mammillary body, almost all the cells expressed Kiss1r on Slc32a1-negative cells. N.B. the wide and strong expression of Kiss1r in PMv that almost every Nissl-stained nucleus contained Kiss1r strongly suggest that those kisspeptin expressing neurons demonstrated in a recent report (1) use autocrine signaling mechanism. Panel **13**: RP3V, rostro-periventricular n. of 3rd ventricle, where Kiss1r is widely expressed in both Slc32a1 positive and negative cells. Panel **14**: examination of RP3V using 4 channels RNAscope LS Multiplex Fluorescent Assay method against mRNA for kisspeptin receptor (Kiss1r, red dots), kisspeptin (Kiss1, green dots), VGAT (Slc32a1, blue dots) and VGLUT2 (Slc17a6, white dots). Six out nine cells (6/9) in this confocal microphotograph were found co-expressing the 4 mRNAs, indicated with quadri-color-arrows. Insets show 4-channel separated screenshots of the cell in the center. The red thin arrows, generated by the Leica Application Suite (LAS) software, indicate the very same coordinates in each channel and the overlay. The three cells which were not Kiss1+ (not Slc32a1+) were all Slc17a6 (VGLUT2) expressing as well as Kiss1r expressing. Panel **15**: low magnification micrograph showing relevant hypothalamic nuclei regarding Kiss1r and Slc32a1 expression. VMN, ventromedial hypothalamic n., Arc, arcuate hypothalamic n., TMv, tuberomammillary n. ventral division, PM, premammillary n. ventral and dorsal divisions, SUM, supramammillary n., MBO, mammillary body. Panel **16**: Kiss1r in ventromedial hypothalamic n. (VMH) is widely and exclusively expressed in no-Slc32a1 cells. In contrast, in the arcuate hypothalamic n. (Arc), the Kiss1r is expressed in both Slc32a1 expressing cells (bicolor arrows) and cells which do not express Slc32a1. Panel **17**: examination of Arc using 4 channels RNAscope LS Multiplex Fluorescent Assay against mRNA for kisspeptin receptor (Kiss1r, red dots), kisspeptin (Kiss1, green dots), VGAT (Slc32a1, blue dots) and VGLUT2 (Slc17a6 white dots). One out of three (1/3) cells in this confocal microphotograph was found co-expressing the mRNAs for Kiss1r, and Slc17a6 Kiss1 expressing but not the Slc32a1 (indicated with tricolor arrows). Insets show 4-channel separated screenshots of the cell in the center. The red thin arrows, generated by the Leica Application Suite (LAS) software, indicate the very same coordinates in each channel. Panel **18**: TMv, tuberomammillary n. ventral division: Kiss1r is expressed in both Slc23a1, and no-Slc32a1 cells. Panel **19**: Kiss1r is strongly expressed in MBO as already shown in panel 12 but weakly expressed in the supramammillary n. (SUM). Panels **20** and **21**: Kiss1r expression in accessory olfactory bulb (AOB) with similar patterns of MOB (see panels 1 and 2), i.e., strongly co-expressed with Slc32a1 (VGAT expressing cells) in periglomerular cells in gl. and granule cells. In the projection layer (mi), the mitral cells which do not express Slc32a, also strongly expressed Kiss1r. The AOB is primarily involved in processing pheromonal signals detected by the vomeronasal organ (VNO) and is critical for mediating social and reproductive behaviors. Panel **22** and **23**: claustrum (CLA) and endopiriform n. (EP), Kiss1r is mainly expressed in no-Slc32a1 neurons. Kiss1r is mainly co-expressed in Slc32a1 (VGAT expressing) neurons in caudoputamen (CP, panel **24**), nucleus accumbens (ACB, panel **25**), nucleus of diagonal band (NDB, panel **26),** bed nucleus of stria terminalis (BST, panel **28**) and reticular thalamic nucleus (Rt, panel **29**, left). Panel **27** shows the mixed expression of Kiss1r, in both Slc32a1postive and negative cells in the lateral preoptic area (LPO). In thalamus, the Kiss1r is widely expressed (panel **29**, right half). In the hippocampal formation, Kiss1r is strongly expressed in the projection neurons (Panels **30, 31, 32, 33** and **34**). Panel **35** and **36**: midbrain superior colliculus (SC) and periaqueductal grey (PAG), Kiss1r mainly expressed in no-Slc32a1 expressing cells. Panel **37** and **38**: pontine parabrachial n. (PB) and locus coeruleus (LC), Kiss1r mainly expressed in no-Slc32a1 expressing cells. Panel **39**: examination of cerebellum using 4 channels RNAscope LS Multiplex Fluorescent Assay against mRNA for kisspeptin (negative and not showing), kisspeptin receptor (Kiss1r, red dots), VGAT (Slc32a1, light blue patches) and VGLUT2 (Slc17a6, white patches). Interposed nucleus was found co-expressing the mRNAs for Kiss1r and Slc17a6 but not the Slc32a1. The multicolor arrows, generated by the Leica Application Suite (LAS) software, indicate the very same coordinates in each channel. Panel **40**: wide expression of Kiss1r in cerebellar granule cell layer (gr). Panel **41**: wide expression of Kiss1r in the brainstem nucleus tractus solitarii (NTS). Panel **42**, **43** and **44**, midbrain ventral tegmental area (VTA), substantia nigra reticulata (SNr), interbrain subthalamic nucleus (STN) and zona incerta (ZI). In the regions 40-44, the Kiss1r is expressed almost exclusively in Slc32a1-negative cells. Panel **45** shows the strong Kiss1r expression in pyramidal cell layer of the piriform cortex (Pir). Panel **46, 47, 49** and **51**: nucleus of lateral olfactory tract (NLOT), cortico-amygdalar nucleus (CoA), basomedial amygdala (BMA) and medial amygdala (MeA): Kiss1r is widely expressed in Slc32a1- negative cells of these basal forebrain structures. Panel **48**: nucleus basalis of Meynert (NBM), a group of cholinergic neurons located in the basal forebrain, widely expressed the Kiss1r. Panel **50**: CEA, central amygdala, Kiss1r is mainly expressed in Slc32a1-postive cells. Scale bars: indicated for each micrograph.

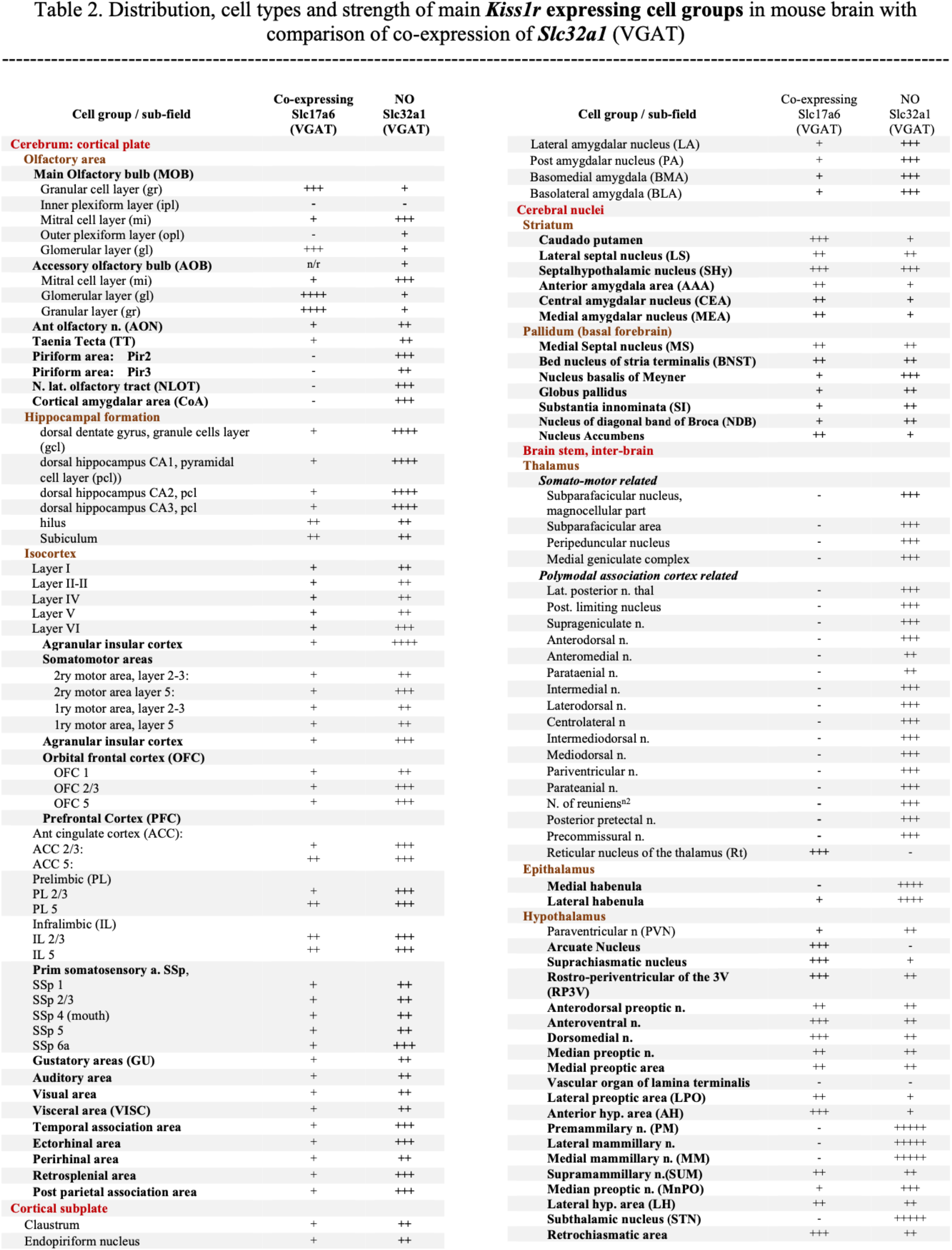
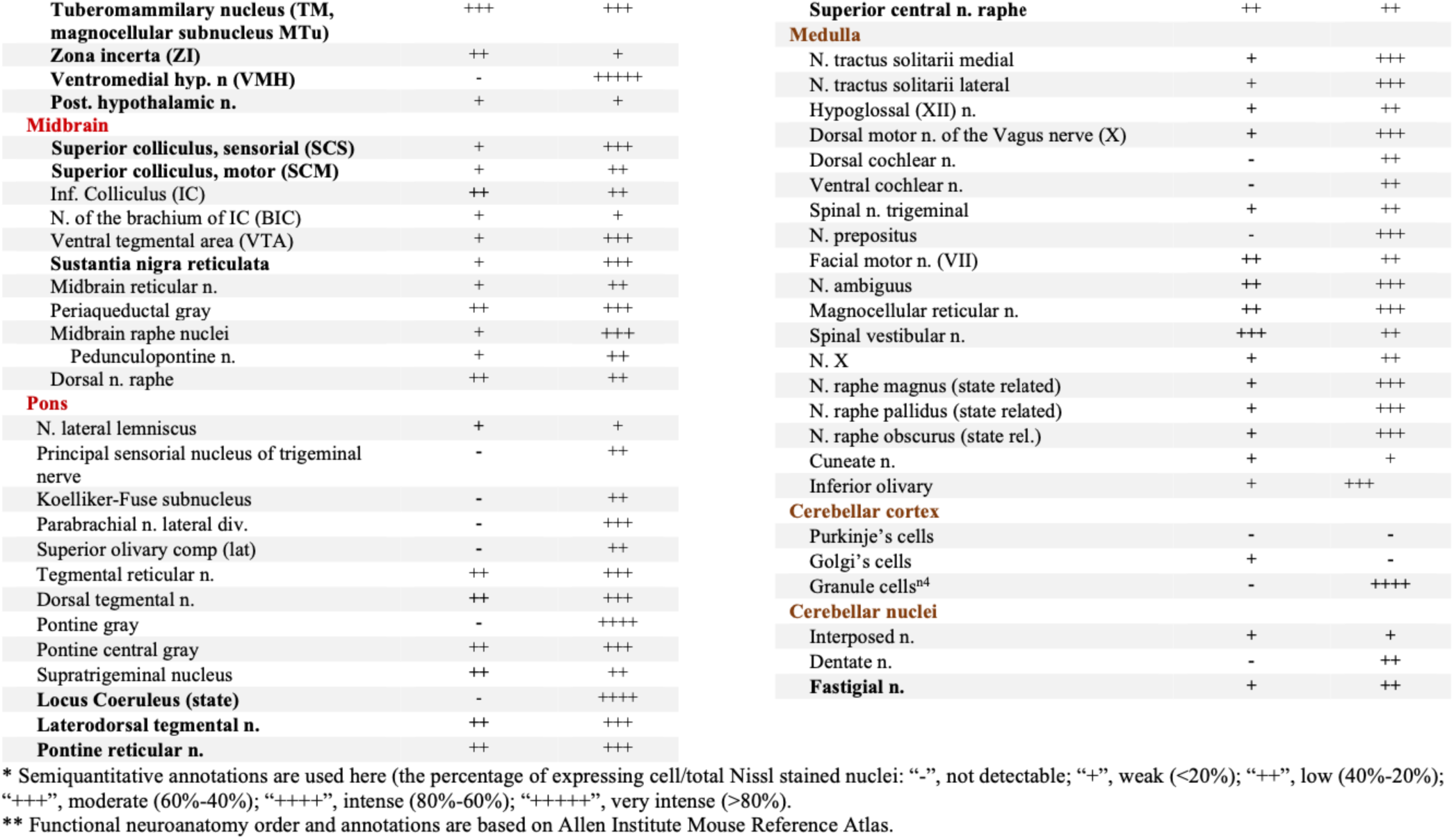

## 4. General discussion and conclusions

This report continues our recent work(1) in which we provided a chemotype of seven *Kiss1*-expressing neuronal populations in the mouse brain. Our current study focuses on the distribution and density of kisspeptin (KP)-immunoreactive (KP-ir) fibers and cell bodies throughout the brains of mice and rats, as well as the distribution of the Kiss1 receptor (*Kiss1r*) in correlation with GABAergic neurons throughout the mouse brain and kisspeptinergic neurons in the hypothalamus.

### 4.1 Prominent KP fiber distribution in sensorial and brain state control centers: ageing and sex steroids

Our anatomical observations regarding the regional distribution of KP-ir fibers and cells in young and aged mice, along with the effects of short- and long-term gonadectomy, underscore the broader roles of KP signaling beyond reproductive functions, potentially including its involvement in sensory processing, behavioral state control, and emotional regulation. Additionally, the lack of a significant age-related decline in KP immunoreactivity - contrary to what is observed in other neuropeptide systems - raises intriguing questions about its role in aging-related diseases such as cancer(68), and neurodegeneration(69), as well as the mechanisms underlying its stability(40).

Aging has been associated with reductions in the density or functionality of various neuropeptide systems, such as neuropeptide Y (NPY), pituitary adenylate cyclase-activating polypeptide (PACAP), and pro-opiomelanocortin (POMC)(70–72) often correlating with cognitive decline, disrupted behavioral states, and changes in physiological homeostasis. With regard to the kisspeptin (KP) system, contrasting effects of aging on KP expression have been documented. It has been reported through immunohistochemistry (IHC) that in aged male and female rats, KNDy-ARC neurons reduce their expression of KP, dynorphin, and neurokinin B peptides(73). In contrast, studies in monkeys and postmortem human tissue reveal that aging correlates with a significant increase in the size of KiSS-1 mRNA-expressing neurons, as well as an increase in the number of labeled cells and autoradiographic grains per neuron(74).

The distribution patterns we report here indicate that many hypothalamic and extrahypothalamic regions are innervated by kisspeptin as strongly as the classical hypothalamic regions known for regulating the hypothalamic-pituitary-gonadal (HPG) axis. These may serve both reproduction-related behavioral, as well as non-reproductive, functions. Indeed, many of these regions are critical for regulating brain states, including telencephalic cortical areas basal forebrain (diagonal band nucleus, accumbens nucleus, septohypothalamic nucleus, medial and lateral septal nuclei), diencephalic structures (thalamic regions which are crucial for cortical activation, arousal and control of awareness such as reuniens nucleus, paraventricular thalamic nucleus anterior(75), hypothalamic median preoptic nucleus, suprachiasmatic nucleus (SCN), complex of the bed nucleus of stria terminalis (BST) and nucleus basalis of Meynert (NBM), midbrain regions (periaqueductal grey, dorsal raphe nucleus, substantia nigra and ventral tegmental area), and metencephalic areas such as the raphe nuclei, reticular formation, and locus coeruleus (see section 3.2 for more focused discussion).

A highlight of this study is the analysis of brains from aged male and female mice gonadectomized early in the lifespan. Notably, the KP fiber density in 22 regions associated with reproductive and non-reproductive functions was found to be similar to, or even increased in, gonadectomized animals compared to controls, suggesting that the KP system remains functional even after reproductive senescence.

KP signaling modulates sexual and emotional brain processing in humans(43). Ogawa et al. found that kisspeptin in the habenula of zebrafish plays a role in fear responses(12, 76), suggesting that it may facilitate the removal of fear, which is essential for maintaining reproductive capabilities under stress. Kisspeptin’s influence on anxiety-related behaviors has also been documented in kisspeptin receptor-deleted male mice, indicating that kisspeptin signaling may play a role in facilitating the activity of anxiogenic neural circuits operative in the elevated plus maze test (EPM)(8).

The interaction between kisspeptin and other neuropeptides was suggested by the work of Stincic et al., in which inputs to PVN and DMH nuclei from presumptive ARC^KP^ and R3PV^KP^ neurons were demonstrated by electrophysiology and optogenetics experiments, suggesting the interaction of the KP system with other neuropeptide systems involved in homeostatic and autonomic regulation(77). Effects of ICV injection of KP-13 suggest that this peptide stimulates the HPA axis, induces hyperthermia, activates motor behavior and causes anxiety in rats(78).

We have reported in this study that a high density of KP fibers is present in areas related to pain sensorial processing, such as the principal sensorial trigeminal nucleus, the gelatinous layer of the trigeminus and in the periaqueductal gray. In the literature, only a few studies have investigated KP’s role in the modulating of the perception of exteroceptive or interoceptive non-reproductive functions. It has been shown that KP-10 activates calcium signaling pathways in cultured dorsal root ganglion cells, suggesting a possible participation in pain modulation(79). Other regions classically involved in sensorial processing have moderate/dense innervation of KP fibers, *i.e.,* superior colliculus for visual processing; the inferior colliculus and superior olive for audition; the parabrachial nucleus and nucleus of the solitary tract (NTS or Sol) for gustatory processing; and the olfactory tubercle, nucleus of the olfactory tract and medial amygdala for olfaction. To our knowledge, the effect of KP on the activity of these regions and its effect on sensorial integration has yet to be examined.

### 4.2 KP receptor distribution suggests potential autocrine, paracrine and neurotransmission modes for KP signaling

Our findings highlight the significant expression of the KP receptor (*Kiss1r*) in various brain regions involved in sensory processing and state control, including the main and accessory olfactory bulbs, septal nuclei, tuberomammillary nuclei, suprachiasmatic nucleus, locus coeruleus, hippocampal pyramidal and granule cell layers, thalamic nuclei, cerebellar granule cell layer, and interposed nucleus. Additionally, *Kiss1r* was expressed in *Kiss1*-expressing hypothalamic cells, indicating a potential autocrine/paracrine role for the peptide, as discussed in detail in Section 3.4.

This expression pattern aligns with previous literature that identifies *Kiss1r* in the hippocampus, particularly in the dentate gyrus—an area critical for learning and memory(30, 80). Activation of *Kiss1r* in the hippocampus has been shown to increase the amplitude of excitatory postsynaptic currents (EPSCs), suggesting a potential mechanism by which kisspeptin may enhance synaptic plasticity(80). In the habenula, prominent expression of *Kiss1* and *Kiss1r* has been reported(26, 30, 31), implicating kisspeptin signaling in emotional regulation as shown in the zebrafish where it has been linked to the inhibition of fear responses(12). Additionally, expressions of Kiss1r have been reported in the amygdala, a region pivotal for processing emotions such as fear and anxiety(26, 31), and the septum, which is involved in reward and reinforcement processes(30). Electrophysiological recordings demonstrate that gonadotropin-releasing hormone (GnRH) septal neurons exhibit robust excitatory responses to kisspeptin application (81). Moreover, the thalamus, known as a relay center for sensory information, also expresses Kiss1r(26, 30)), while reports indicate its presence in medullary and pontine brainstem regions(30, 82). Kiss1r expression has been reported in the medial amygdala (MeApd) of male rats, contributing to the facilitation of erectile function(61). Finally, the striatum and cortical regions have been also shown to express Kiss1r(26, 82) however we found no studies investigating the potential role of KP signaling in these regions. Collectively, these findings underscore the complex and diverse roles of Kiss1r throughout the brain, suggesting that the kisspeptin signaling pathway may be instrumental in modulating a wide array of sensory and emotional processes.

### 4.3 Controversies and some consideration

Rather dramatic down-regulation in mRNA for kisspeptin (*Kiss1)* expression in the RP3V and up- regulation in the Arc region, in term of dendritic mRNA expression, after acute gonadectomy and a generally stronger Kiss1 expression in female mice vs males was recently reported (1), in agreement with most of the previous reports characterizing KP mRNA expression in this region in both intact and gonadactomized male and female rodents(83–86). However, altered KP expression at the level of the nerve terminal is more difficult to discern, perhaps because immunohistochemical terminal field appearance in a given brain area is determined by the composite projections from multiple KP cell groups. In SI Figure 2, we analyzed the experiments from Allen Mouse Brain Connectivity Atlas open source to demonstrate the reciprocal and convergent Kiss1 innervations in mouse brain of KP^RP3V^ and KP^Arc^ and in SI Figure 3 we presented this notion in diagram. It is our hope that future investigations will be directed towards more granular analysis of the individual contributions of each KP neuronal population to the nerve terminals within given brain areas.

#### Contribution statement

Conceptualization: LZ, VSH, MAZ, RPM, LEE; data curation: LZ, VSH, MAZ, ORHP; investigation: LZ, VSH, MAZ, ORHP, RHG; methodology: LZ, VSH, RHG; project administration: LZ, VH, LEE; resources: LZ, LEE, RPM, ICA, RHG; visualization LZ, VSH; writing – original draft: LZ, VH, LEE; review & editing: all the co-authors.

## Acknowledgement

The research leading to these results received funding from the National Autonomous University of Mexico, under Grant Agreement UNAM-PAPIIT-IG200121(LZ), and UNAM-PAPIIT-IA202724 (ORHP); Mexican Nacional Council for Humanity, Science and Technology (CONAHCYT), under Grant Agreement CF-2023-243 (LZ), NIMH-IR, NIH, under Grant Agreement MH002386 (LEE). The investigators of this study have been supported by the following fellowships: sabbatical year fellowship from the PASPA program of the Dirección General de Personal Académico (DGAPA) of the Universidad Nacional Autónoma de México (UNAM) (LZ, VSH); Mexican CONAHCYT sabbatical fellowship (LZ) and Fulbright-García Robles Fellowship (VSH), for sabbatical research year in LEE’s laboratory of NIMH, NIH; DGAPA-UNAM PREI program for a sabbatical research stay (RPM) in LZ’s lab in UNAM, Mexico. We thank Charles Gerfen, NIMH for his support and discussions about this study. We thank Mr. Jonathan Kuo and colleagues of the NIMH-IRP Systems Neuroscience Imaging Resource (SNIR, Director Ted Usdin) and Ms. Ivonne Sanchez Cervantes of the Microscopy Unit of the School of Medicine, UNAM for technical assistance.

---

**Supplementary Figure 1.**
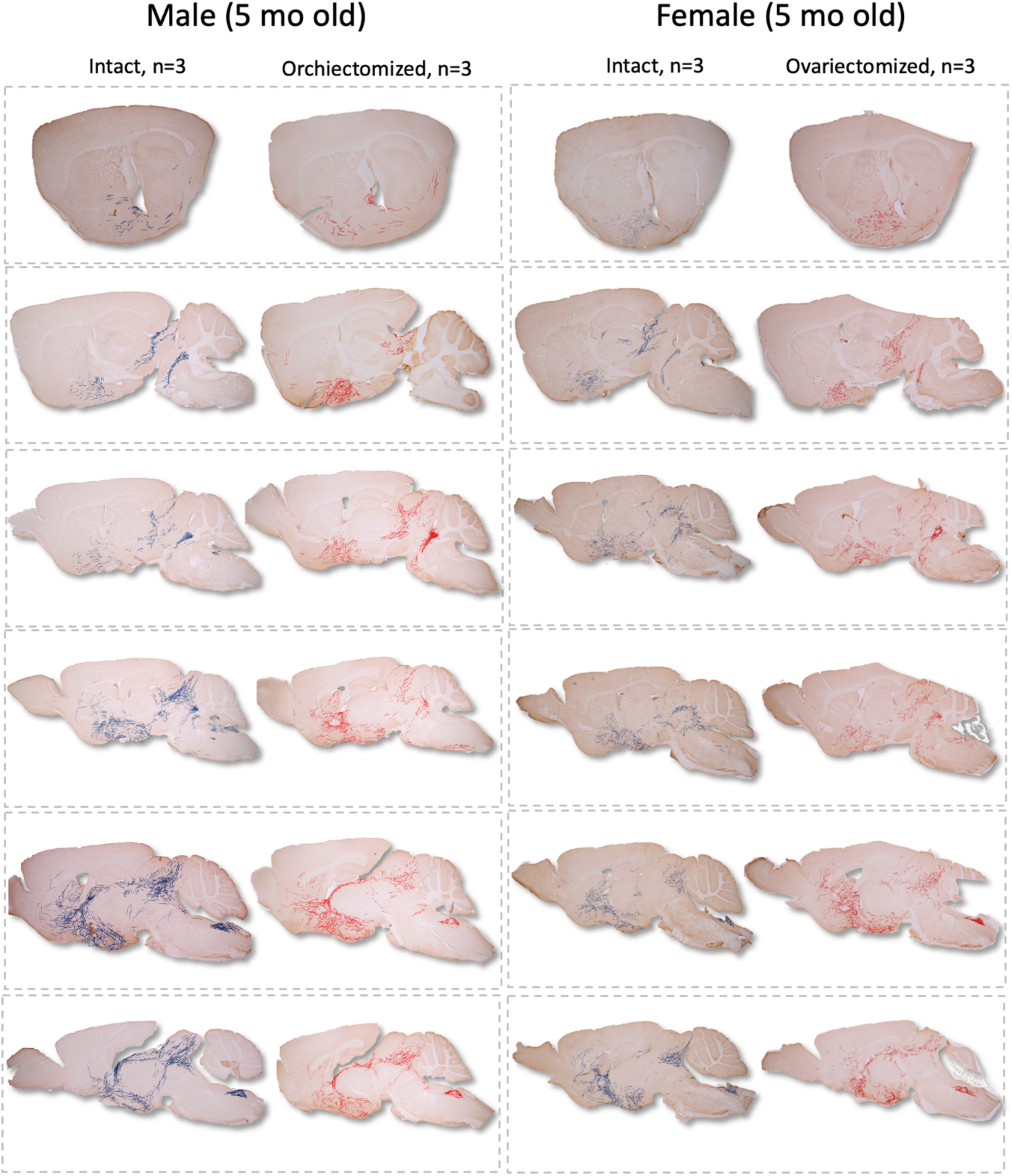
Comparison of cumulative (summed) fiber density charting of three young adult animals from each of the 4 experimental groups (male/female and intact/gonadectomized, N=12) represented in six different parasagittal levels. Hand-made drawings under microscope observations highlight the fiber distributions abstracted from the 3 subjects of the same group. Lines (control: blue, GNX, red) representing the KP-ir fibers which are "exaggerated" in term of thickness, in relation to the section area, for visualization sake.

**Supplementary Figure 2.**
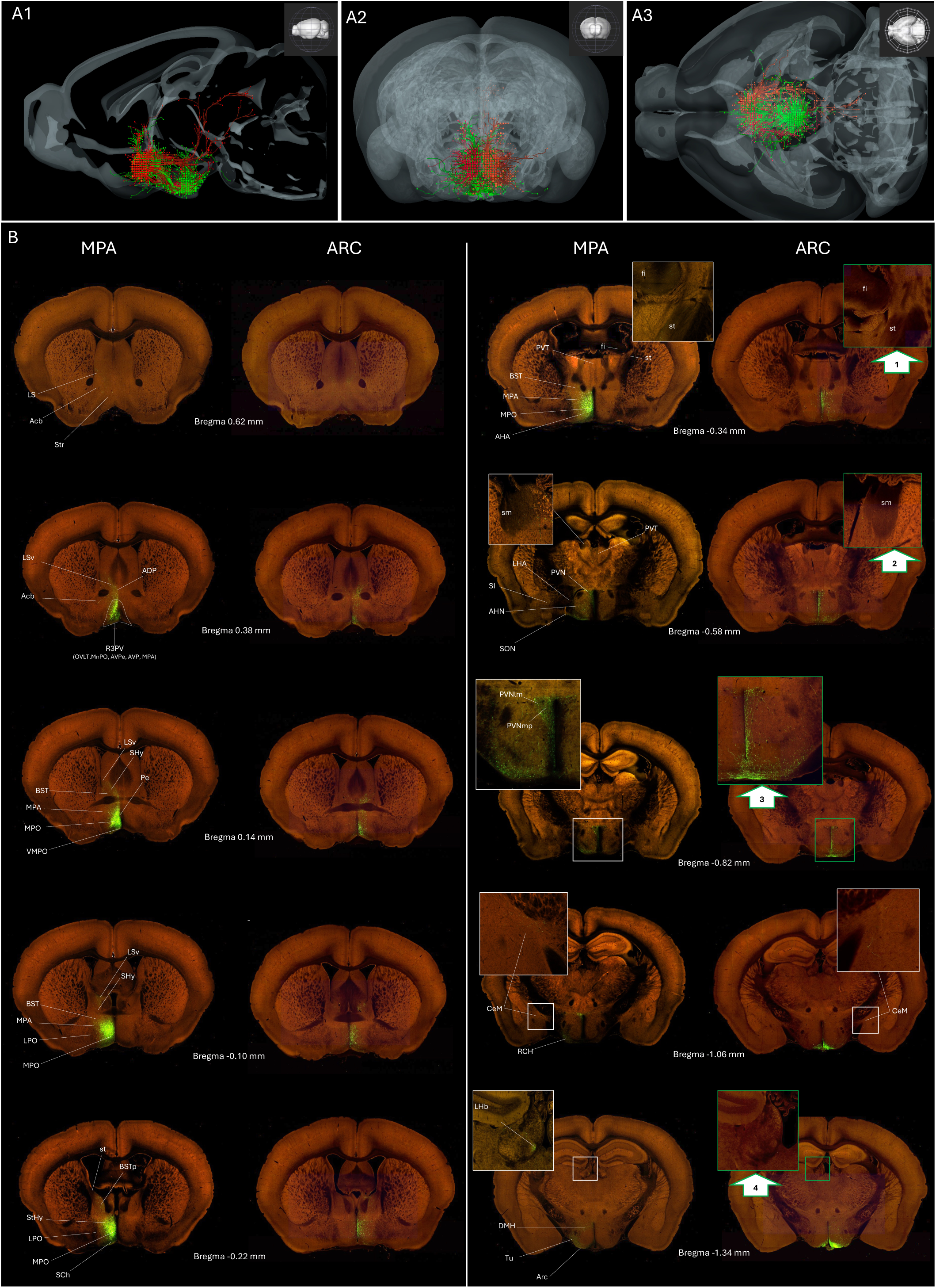
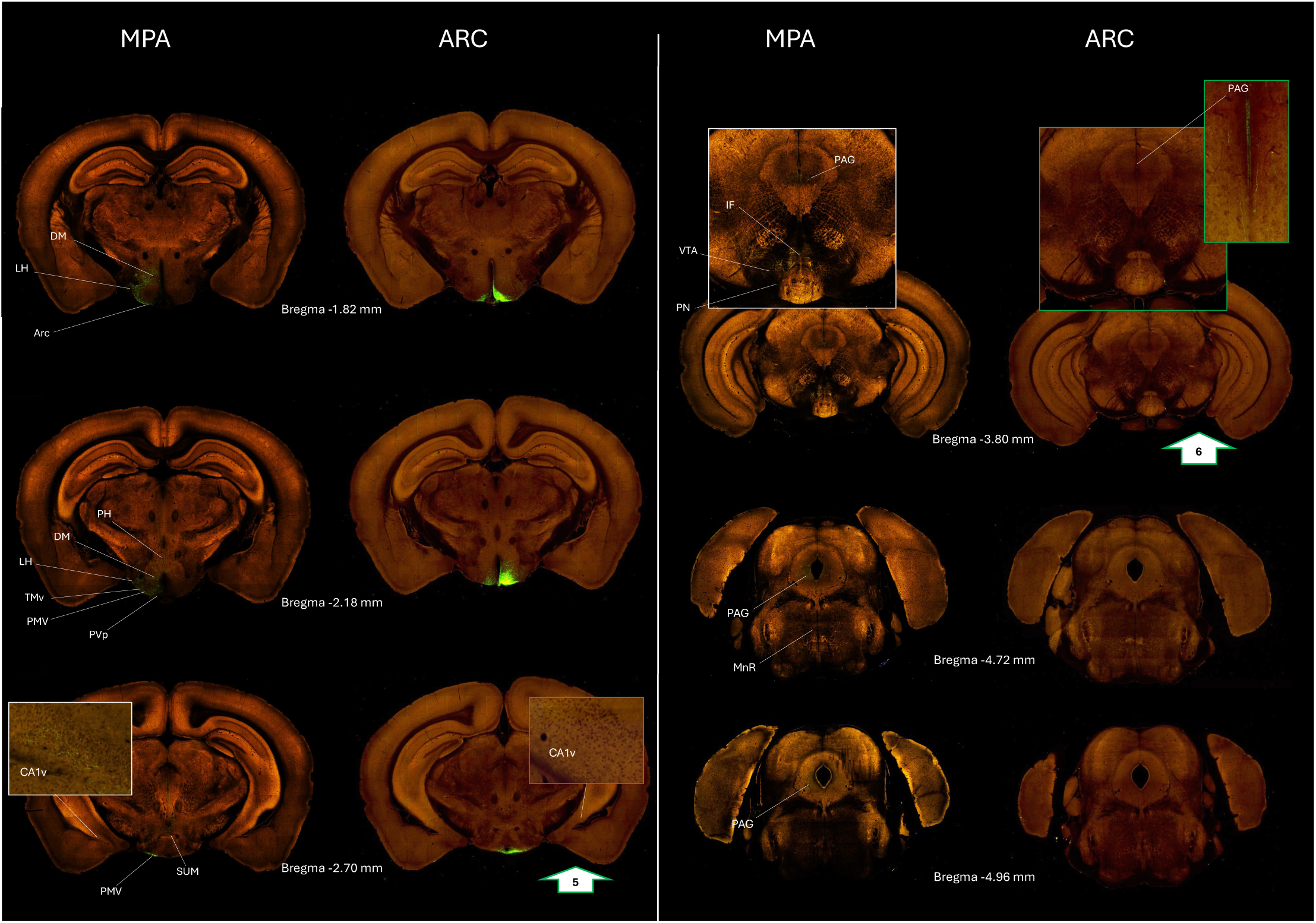
**Projection targets of R3PV or Arcuate Kiss1-Cre neurons transfected with AAV**. The data from two Kiss1-Cre mice injected in the arcuate nucleus (Arc, experiments 232311236 and 232310521) and two Kiss1-Cre mice injected in the rostral periventricular region (R3PV, experiments 299247009 and 301989585) injected with Cre-dependent tracer from the displayed in Allen Mouse Brain Connectivity Atlas this figure. Sagittal (A1), coronal (A2) and horizontal (A3) views of a 3-D representation with 3-D brain explorer of the innervation patterns of the two major populations of Kiss1 neurons. The VGAT-expressing R3PV neuronal population and its projections is represented in red, and the VGLUT2-expressing Arc population and its projections is represented in green. The different views show a reciprocal dense innervation between the Arc and the RP3V, additionally it is clear that the majority of regions in the striatum, septum, BNST, thalamus, hypothalamus, and amygdala, receive overlapping innervation from both regions. The ponto-mecencephalic region is mainly innervated by the R3PV. (B) Coronal sections at 17 rostro-caudal Bregma coordinates were selected from the animal that showed the best labelling for every level, for each level the left section from a R3PV injected mice was paired with a section from the Arc-injected mice. Some regions of interest are labelled in the RP3V sections, green-numbered arrows in the Arc sections indicate regions where there were differences in the innervation patterns between RP3V and Arc injected mice. Arrow #1: Some fibers were observed in the main stria terminalis and fimbria in the RP3V injected mice, but not in the Arc injected. Arrow #2: some fibers were observed in the stria medularis in the RP3v but not in the Arc injected mice. Arrow 3: The pattern of innervation of the paraventricular hypothalamic nucleus (PVN) is different between these experiments, the labelled axons of the mice injected in the RP3V innervate the magnocellular and parvocellular division, and the Arc injected neurons innervate preferentially the parvocellular region of the PVN. Also notice at this level and the following (Bregma -1.06) that for both injection targets, some axons travel laterally to reach the CeM. Arrow #4: the mice injected in the RP3V show innervation of the medial part of the lateral habenula (LHbm) specially the rostral part, thes fibers appear to travel via the stria medullaris (see arrow 2). No fibers in LHb were found in the Arc experiments. Arrow #5: A plexus of labelled fibers can be observed in the CA1 pyramidal layer of the ventral hippocampus, these fibers originate in the RP3V and travel trough the stria terminalis (see arrow 1). No fibers were found at this location in the Arc injected mice. Arrow #6: Fibers from the RP3V reach the mecencephalic region and innervate the periaquieductal gray region (PAG), the interfascicular nucleus (IF), the ventral tegmental area (VTA) and the median raphe (MnR). The mice injected in the Arc, show sparse axons and mainly innervate the PAG.

**Supplementary Figure 3.**
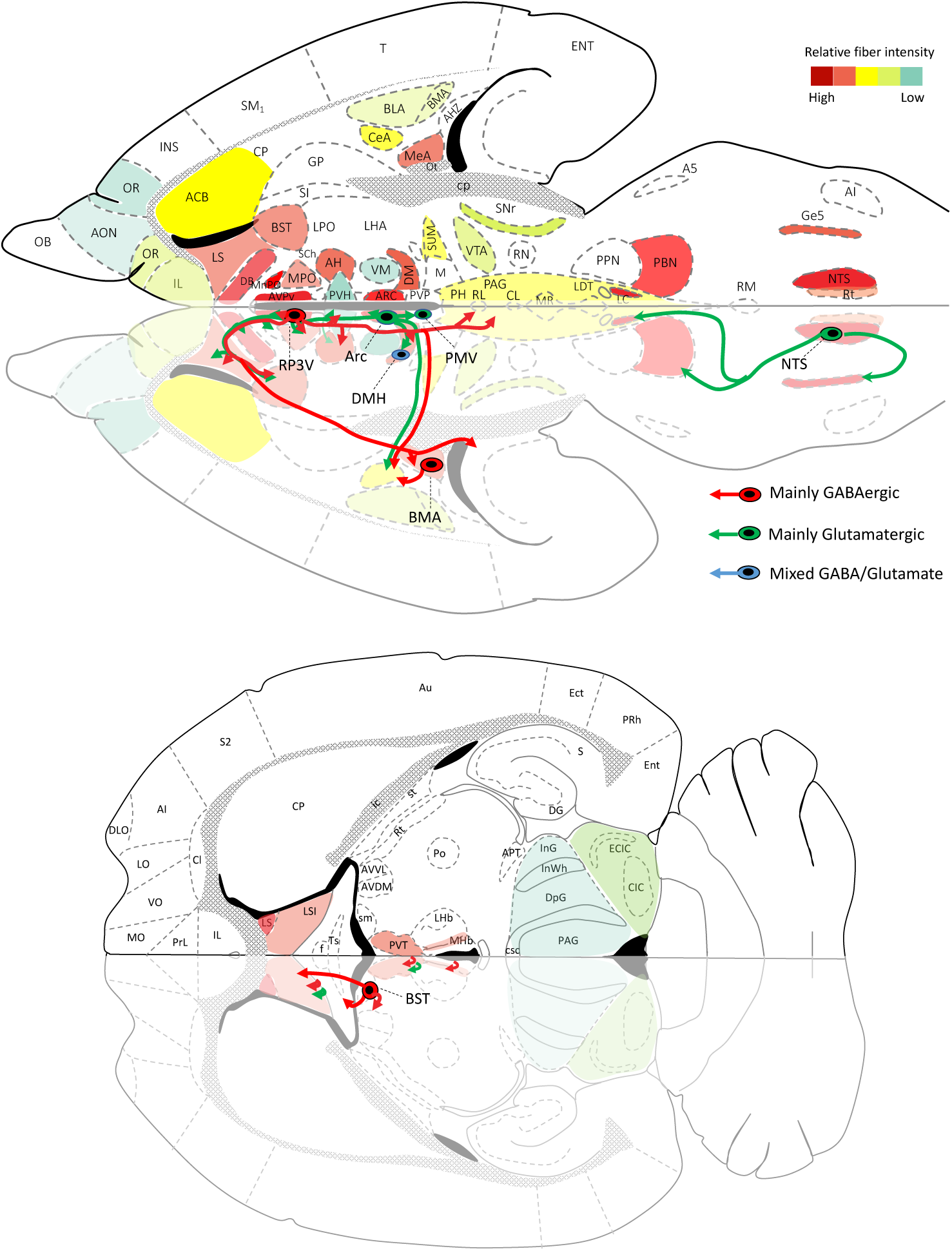
Kisspeptin fiber density Kisspeptin pathways to regions involved in brain state control, as well as motor and sensorial processing. The density of kisspeptin observed through the brain of the "ideal mouse" i.e. the maximum density found in males or females through all gonadal states is represented as a relative fiber density, color coded in two horizontal levels of flatmap where some of the most innervated regions related to brain state, sensorial processing and motor integration were found. The bottom part of the drawing represent the pathways that we identified using analysis of several consecutive immunoreacted slices in different planes, as well as data obtained from four experiments published in the Allen brain atlas. the seven populations described in this report are labelled and shown in red if they are mainly Slc32A1 (VGAT) expressing or in green inf they are mainly Slc17a6 (VGLUT2) expressing. The population of the DMH was hosted GABAergic and Glutamatergic neurons and is coded in blue.

## References

1. Hernández VS, Zetter MA, Hernández-Pérez O, Hernández-González R, Camacho-Arroyo I, Millar RP, Eiden LE, Zhang L. A comprehensive chemotyping and gonadal regulation of seven kisspeptinergic neuronal populations in the mouse brain. bioRxiv. 20242024.07.23.604881.

2. Lee JH, Miele ME, Hicks DJ, Phillips KK, Trent JM, Weissman BE, Welch DR. KiSS-1, a novel human malignant melanoma metastasis-suppressor gene. J Natl Cancer Inst. 1996; 88(23): 1731–7.

3. Pinilla L, Aguilar E, Dieguez C, Millar RP, Tena-Sempere M. Kisspeptins and reproduction: physiological roles and regulatory mechanisms. Physiol Rev. 2012; 92(3): 1235–316.

4. Clarke H, Dhillo WS, Jayasena CN. Comprehensive Review on Kisspeptin and Its Role in Reproductive Disorders. Endocrinol Metab (Seoul). 2015; 30(2): 124–41.

5. Mills EGA, O’Byrne KT, Comninos AN. Kisspeptin as a Behavioral Hormone. Semin Reprod Med. 2019; 37(2): 56–63.

6. Mills EG, Yang L, Abbara A, Dhillo WS, Comninos AN. Current Perspectives on Kisspeptins Role in Behaviour. Front Endocrinol (Lausanne). 2022; 13 928143.

7. Stephens SBZ, Kauffman AS. Regulation and Possible Functions of Kisspeptin in the Medial Amygdala. Front Endocrinol (Lausanne). 2017; 8191.

8. Delmas S, Porteous R, Bergin DH, Herbison AE. Altered aspects of anxiety-related behavior in kisspeptin receptor-deleted male mice. Sci Rep. 2018; 8(1): 2794.

9. Ebrahimi Khonacha S, Janahmadi M, Motamedi F. Kisspeptin-13 Improves Spatial Memory Consolidation and Retrieval against Amyloid-beta Pathology. Iran J Pharm Res. 2019; 18(Suppl1): 169–81.

10. Abdul Satar NM, Ogawa S, Parhar IS. Kisspeptin-1 regulates forebrain dopaminergic neurons in the zebrafish. Sci Rep. 2020; 10(1): 19361.

11. Sivalingam M, Ogawa S, Parhar IS. Habenula kisspeptin retrieves morphine impaired fear memory in zebrafish. Sci Rep. 2020; 10(1): 19569.

12. Ogawa S, Nathan FM, Parhar IS. Habenular kisspeptin modulates fear in the zebrafish. Proc Natl Acad Sci U S A. 2014; 111(10): 3841–6.

13. Liu X, Herbison AE. Kisspeptin Regulation of Neuronal Activity throughout the Central Nervous System. Endocrinol Metab (Seoul). 2016; 31(2): 193–205.

14. Clarkson J, d’Anglemont de Tassigny X, Colledge WH, Caraty A, Herbison AE. Distribution of kisspeptin neurones in the adult female mouse brain. J Neuroendocrinol. 2009; 21(8): 673–82.

15. Muir AI, Chamberlain L, Elshourbagy NA, Michalovich D, Moore DJ, Calamari A, Szekeres PG, Sarau HM, Chambers JK, Murdock P, Steplewski K, Shabon U, Miller JE, Middleton SE, Darker JG, Larminie CG, Wilson S, Bergsma DJ, Emson P, Faull R, Philpott KL, Harrison DC. AXOR12, a novel human G protein-coupled receptor, activated by the peptide KiSS-1. J Biol Chem. 2001; 276(31): 28969–75.

16. Lehman MN, Hileman SM, Goodman RL. Neuroanatomy of the kisspeptin signaling system in mammals: comparative and developmental aspects. Adv Exp Med Biol. 2013; 784 27–62.

17. Mikkelsen JD, Simonneaux V. The neuroanatomy of the kisspeptin system in the mammalian brain. Peptides. 2009; 30(1): 26–33.

18. Lehman MN, Coolen LM, Steiner RA, Neal-Perry G, Wang L, Moenter SM, Moore AM, Goodman RL, Hwa-Yeo S, Padilla SL, Kauffman AS, Garcia J, Kelly MJ, Clarkson J, Radovick S, Babwah AV, Leon S, Tena-Sempere M, Comninos A, Seminara S, Dhillo WS, Levine J, Terasawa E, Negron A, Herbison AE. The 3(rd) World Conference on Kisspeptin, "Kisspeptin 2017: Brain and Beyond":Unresolved questions, challenges and future directions for the field. J Neuroendocrinol. 2018e12600.

19. Zhang L, Hernandez VS. Synaptic innervation to rat hippocampus by vasopressin-immuno- positive fibres from the hypothalamic supraoptic and paraventricular nuclei. Neuroscience. 2013; 228 139–62.

20. Swanson LW. Brain Architecture: Understanding the Basic Plan Oxford, UK: Oxford University Press, 2011.

21. Lee SH, Dan Y. Neuromodulation of brain states. Neuron. 2012; 76(1): 209–22.

22. Zagha E, McCormick DA. Neural control of brain state. Curr Opin Neurobiol. 2014; 29 178–86.

23. Rosazza C, Minati L. Resting-state brain networks: literature review and clinical applications. Neurol Sci. 2011; 32(5): 773–85.

24. McCormick DA, Nestvogel DB, He BJ. Neuromodulation of Brain State and Behavior. Annu Rev Neurosci. 2020; 43 391–415.

25. Berridge CW, Waterhouse BD. The locus coeruleus–noradrenergic system: modulation of behavioral state and state-dependent cognitive processes. Brain research reviews. 2003; 42(1): 33–84.

26. Lee DK, Nguyen T, O’Neill GP, Cheng R, Liu Y, Howard AD, Coulombe N, Tan CP, Tang-Nguyen AT, George SR, O’Dowd BF. Discovery of a receptor related to the galanin receptors. FEBS Lett. 1999; 446(1): 103–7.

27. Franssen D, Tena-Sempere M. The kisspeptin receptor: A key G-protein-coupled receptor in the control of the reproductive axis. Best Pract Res Clin Endocrinol Metab. 2018; 32(2): 107–23.

28. Irwig MS, Fraley GS, Smith JT, Acohido BV, Popa SM, Cunningham MJ, Gottsch ML, Clifton DK, Steiner RA. Kisspeptin activation of gonadotropin releasing hormone neurons and regulation of KiSS-1 mRNA in the male rat. Neuroendocrinology. 2004; 80(4): 264–72.

29. Ringel MD, Hardy E, Bernet VJ, Burch HB, Schuppert F, Burman KD, Saji M. Metastin receptor is overexpressed in papillary thyroid cancer and activates MAP kinase in thyroid cancer cells. J Clin Endocrinol Metab. 2002; 87(5): 2399.

30. Herbison AE, de Tassigny X, Doran J, Colledge WH. Distribution and postnatal development of Gpr54 gene expression in mouse brain and gonadotropin-releasing hormone neurons. Endocrinology. 2010; 151(1): 312–21.

31. Higo S, Honda S, Iijima N, Ozawa H. Mapping of Kisspeptin Receptor mRNA in the Whole Rat Brain and its Co-Localisation with Oxytocin in the Paraventricular Nucleus. J Neuroendocrinol. 2016; 28(4).

32. Stout Steele M, Bennett RA. Clinical Technique: Dorsal Ovariectomy in Rodents. Journal of Exotic Pet Medicine. 2011; 20(3): 222–6.

33. Sophocleous A, Idris AI. Ovariectomy/Orchiectomy in Rodents. In: Idris AI, ed. Bone Research Protocols. New York, NY: Springer New York 2019: 261-7.

34. Dhillo WS, Chaudhri OB, Patterson M, Thompson EL, Murphy KG, Badman MK, McGowan BM, Amber V, Patel S, Ghatei MA, Bloom SR. Kisspeptin-54 stimulates the hypothalamic-pituitary gonadal axis in human males. J Clin Endocrinol Metab. 2005; 90(12): 6609–15.

35. Paxinos G, Franklin KBJ. Paxinos and Franklin’s the Mouse Brain in Stereotaxic Coordinates Elsevier Science, 2012.

36. Paxinos G, Watson C. The Rat Brain in Stereotaxic Coordinates Elsevier Science, 2007.

37. Zhang L, Hernandez VS, Gerfen CR, Jiang SZ, Zavala L, Barrio RA, Eiden LE. Behavioral role of PACAP signaling reflects its selective distribution in glutamatergic and GABAergic neuronal subpopulations. Elife. 2021; 10.

38. Zhang L, Hernandez VS, Zetter MA, Eiden LE. VGLUT-VGAT expression delineates functionally specialised populations of vasopressin-containing neurones including a glutamatergic perforant path- projecting cell group to the hippocampus in rat and mouse brain. J Neuroendocrinol. 2020; 32(4): e12831.

39. Brailoiu GC, Dun SL, Ohsawa M, Yin D, Yang J, Chang JK, Brailoiu E, Dun NJ. KiSS-1 expression and metastin-like immunoreactivity in the rat brain. J Comp Neurol. 2005; 481(3): 314–29.

40. Ketterson ED, Nolan V, Jr. Adaptation, Exaptation, and Constraint: A Hormonal Perspective. Am Nat. 1999; 154(S1): S4–S25.

41. Risold PY, Swanson LW. Connections of the rat lateral septal complex. Brain research Brain research reviews. 1997; 24(2-3): 115–95.

42. Rizzi-Wise CA, Wang DV. Putting Together Pieces of the Lateral Septum: Multifaceted Functions and Its Neural Pathways. eNeuro. 2021; 8(6).

43. Comninos AN, Wall MB, Demetriou L, Shah AJ, Clarke SA, Narayanaswamy S, Nesbitt A, Izzi- Engbeaya C, Prague JK, Abbara A, Ratnasabapathy R, Salem V, Nijher GM, Jayasena CN, Tanner M, Bassett P, Mehta A, Rabiner EA, Honigsperger C, Silva MR, Brandtzaeg OK, Lundanes E, Wilson SR, Brown RC, Thomas SA, Bloom SR, Dhillo WS. Kisspeptin modulates sexual and emotional brain processing in humans. J Clin Invest. 2017; 127(2): 709–19.

44. Sheehan TP, Chambers RA, Russell DS. Regulation of affect by the lateral septum: implications for neuropsychiatry. Brain research Brain research reviews. 2004; 46(1): 71–117.

45. Salib M, Joshi A, Katona L, Howarth M, Micklem BR, Somogyi P, Viney TJ. GABAergic Medial Septal Neurons with Low-Rhythmic Firing Innervating the Dentate Gyrus and Hippocampal Area CA3. J Neurosci. 2019; 39(23): 4527–49.

46. Unal G, Crump MG, Viney TJ, Eltes T, Katona L, Klausberger T, Somogyi P. Spatio-temporal specialization of GABAergic septo-hippocampal neurons for rhythmic network activity. Brain Struct Funct. 2018; 223(5): 2409–32.

47. Buzsaki G. Theta oscillations in the hippocampus. Neuron. 2002; 33(3): 325–40.

48. Espinosa N, Caneo M, Alonso A, Moran C, Fuentealba P. Optogenetic stimulation of septal somatostatin neurons disrupts locomotory behavior and regulates hippocampus cholinergic theta oscillations. bioRxiv. 20222022.09.30.510330.

49. Shen L, Zhang GW, Tao C, Seo MB, Zhang NK, Huang JJ, Zhang LI, Tao HW. A bottom-up reward pathway mediated by somatostatin neurons in the medial septum complex underlying appetitive learning. Nat Commun. 2022; 13(1): 1194.

50. Srividya R, Mallick HN, Kumar VM. The changes in thermal preference, sleep-wakefulness, body temperature and locomotor activity in the rats with medial septal lesion. Behav Brain Res. 2005; 164(2): 147–55.

51. Tsanov M. Speed and Oscillations: Medial Septum Integration of Attention and Navigation. Front Syst Neurosci. 2017; 1167.

52. Joshi A, Salib M, Viney TJ, Dupret D, Somogyi P. Behavior-Dependent Activity and Synaptic Organization of Septo-hippocampal GABAergic Neurons Selectively Targeting the Hippocampal CA3 Area. Neuron. 2017; 96(6): 1342–57 e5.

53. Liu AKL, Lim EJ, Ahmed I, Chang RC, Pearce RKB, Gentleman SM. Review: Revisiting the human cholinergic nucleus of the diagonal band of Broca. Neuropathol Appl Neurobiol. 2018; 44(7): 647–62.

54. Dudek SM, Alexander GM, Farris S. Rediscovering area CA2: unique properties and functions. Nature reviews Neuroscience. 2016; 17(2): 89–102.

55. Semba K, Fibiger HC. Organization of central cholinergic systems. Prog Brain Res. 1989; 79 37–63.

56. Floresco SB. The nucleus accumbens: an interface between cognition, emotion, and action. Annu Rev Psychol. 2015; 66 25–52.

57. Riaz S, Puveendrakumaran P, Khan D, Yoon S, Hamel L, Ito R. Prelimbic and infralimbic cortical inactivations attenuate contextually driven discriminative responding for reward. Sci Rep. 2019; 9(1): 3982.

58. Capuzzo G, Floresco SB. Prelimbic and Infralimbic Prefrontal Regulation of Active and Inhibitory Avoidance and Reward-Seeking. J Neurosci. 2020; 40(24): 4773–87.

59. de Lima MAX, Baldo MVC, Oliveira FA, Canteras NS. The anterior cingulate cortex and its role in controlling contextual fear memory to predatory threats. Elife. 2022; 11.

60. Kataoka N, Shima Y, Nakajima K, Nakamura K. A central master driver of psychosocial stress responses in the rat. Science. 2020; 367(6482): 1105-12.

61. Gresham R, Li S, Adekunbi DA, Hu M, Li XF, O’Byrne KT. Kisspeptin in the medial amygdala and sexual behavior in male rats. Neurosci Lett. 2016; 627 13–7.

62. Pineda R, Plaisier F, Millar RP, Ludwig M. Amygdala Kisspeptin Neurons: Putative Mediators of Olfactory Control of the Gonadotropic Axis. Neuroendocrinology. 2017; 104(3): 223–38.

63. Kim J, Semaan SJ, Clifton DK, Steiner RA, Dhamija S, Kauffman AS. Regulation of Kiss1 expression by sex steroids in the amygdala of the rat and mouse. Endocrinology. 2011; 152(5): 2020–30.

64. Dubois SL, Acosta-Martinez M, DeJoseph MR, Wolfe A, Radovick S, Boehm U, Urban JH, Levine JE. Positive, but not negative feedback actions of estradiol in adult female mice require estrogen receptor alpha in kisspeptin neurons. Endocrinology. 2015; 156(3): 1111–20.

65. Rance NE. Menopause and the human hypothalamus: evidence for the role of kisspeptin/neurokinin B neurons in the regulation of estrogen negative feedback. Peptides. 2009; 30(1): 111–22.

66. Starrett JR, Moenter SM. Hypothalamic kisspeptin neurons as potential mediators of estradiol negative and positive feedback. Peptides. 2023; 163 170963.

67. Goodman RL, Moore AM, Onslow K, Hileman SM, Hardy SL, Bowdridge EC, Walters BA, Agus S, Griesgraber MJ, Aerts EG, Lehman MN, Coolen LM. Lesions of KNDy and Kiss1R Neurons in the Arcuate Nucleus Produce Different Effects on LH Pulse Patterns in Female Sheep. Endocrinology. 2023; 164(11).

68. Kim TH, Yoon JH, Cho SG. Kisspeptin Promotes Glioblastoma Cell Invasiveness Via the Gq-PLC- PKC Pathway. Anticancer Res. 2020; 40(1): 213–20.

69. Sinen O, Sinen AG, Derin N, Aslan MA. Nasal application of kisspeptin-54 mitigates motor deficits by reducing nigrostriatal dopamine loss in hemiparkinsonian rats. Behav Brain Res. 2024; 468 115035.

70. Klinger K, Del Angel M, Caliskan G, Stork O. Increasing NPYergic transmission in the hippocampus rescues aging-related deficits of long-term potentiation in the mouse dentate gyrus. Front Aging Neurosci. 2023; 15 1283581.

71. Reglodi D, Atlasz T, Szabo E, Jungling A, Tamas A, Juhasz T, Fulop BD, Bardosi A. PACAP deficiency as a model of aging. Geroscience. 2018; 40(5-6): 437–52.

72. Kappeler L, Gourdji D, Zizzari P, Bluet-Pajot MT, Epelbaum J. Age-associated changes in hypothalamic and pituitary neuroendocrine gene expression in the rat. J Neuroendocrinol. 2003; 15(6): 592–601.

73. Kunimura Y, Iwata K, Ishigami A, Ozawa H. Age-related alterations in hypothalamic kisspeptin, neurokinin B, and dynorphin neurons and in pulsatile LH release in female and male rats. Neurobiol Aging. 2017; 50 30–8.

74. Rometo AM, Krajewski SJ, Voytko ML, Rance NE. Hypertrophy and increased kisspeptin gene expression in the hypothalamic infundibular nucleus of postmenopausal women and ovariectomized monkeys. J Clin Endocrinol Metab. 2007; 92(7): 2744–50.

75. Biesbroek JM, Verhagen MG, van der Stigchel S, Biessels GJ. When the central integrator disintegrates: A review of the role of the thalamus in cognition and dementia. Alzheimers Dement. 2024; 20(3): 2209–22.

76. Ogawa S, Parhar IS. Functions of habenula in reproduction and socio-reproductive behaviours. Front Neuroendocrinol. 2022; 64 100964.

77. Stincic TL, Qiu J, Connors AM, Kelly MJ, Ronnekleiv OK. Arcuate and Preoptic Kisspeptin Neurons Exhibit Differential Projections to Hypothalamic Nuclei and Exert Opposite Postsynaptic Effects on Hypothalamic Paraventricular and Dorsomedial Nuclei in the Female Mouse. eNeuro. 2021; 8(4).

78. Csabafi K, Jaszberenyi M, Bagosi Z, Liptak N, Telegdy G. Effects of kisspeptin-13 on the hypothalamic-pituitary-adrenal axis, thermoregulation, anxiety and locomotor activity in rats. Behav Brain Res. 2013; 241 56–61.

79. Kelestimur H, Bulut F, Canpolat S, Ozcan M, Ayar A. Kisspeptin leads to calcium signaling in cultured rat dorsal root ganglion neurons. Gen Physiol Biophys. 2021; 40(2): 155–60.

80. Arai AC. The role of kisspeptin and GPR54 in the hippocampus. Peptides. 2009; 30(1): 16–25.

81. Dumalska I, Wu M, Morozova E, Liu R, van den Pol A, Alreja M. Excitatory effects of the puberty- initiating peptide kisspeptin and group I metabotropic glutamate receptor agonists differentiate two distinct subpopulations of gonadotropin-releasing hormone neurons. J Neurosci. 2008; 28(32): 8003–13.

82. Mattam U, Talari NK, Thiriveedi VR, Fareed M, Velmurugan S, Mahadev K, Sepuri NBV. Aging reduces kisspeptin receptor (GPR54) expression levels in the hypothalamus and extra-hypothalamic brain regions. Exp Ther Med. 2021; 22(3): 1019.

83. Smith JT, Cunningham MJ, Rissman EF, Clifton DK, Steiner RA. Regulation of Kiss1 gene expression in the brain of the female mouse. Endocrinology. 2005; 146(9): 3686–92.

84. Smith JT, Dungan HM, Stoll EA, Gottsch ML, Braun RE, Eacker SM, Clifton DK, Steiner RA. Differential regulation of KiSS-1 mRNA expression by sex steroids in the brain of the male mouse. Endocrinology. 2005; 146(7): 2976–84.

85. Kauffman AS. Gonadal and nongonadal regulation of sex differences in hypothalamic Kiss1 neurones. J Neuroendocrinol. 2010; 22(7): 682–91.

86. Takumi K, Iijima N, Iwata K, Higo S, Ozawa H. The effects of gonadal steroid manipulation on the expression of Kiss1 mRNA in rat arcuate nucleus during postnatal development. J Physiol Sci. 2012; 62(6): 453–60.

